# Rif-seq reveals *Caulobacter crescentus* mRNA decay is globally coordinated with transcription and translation

**DOI:** 10.1101/2025.04.22.648521

**Authors:** H. Kulathungage, J.R. Aretakis, K. Vaidya, N. Al-Husini, N.S. Muthunayake, S. Kim, J.M. Schrader

## Abstract

While transcription and translation have been shown to be coordinated with mRNA decay across various single-gene studies, their global coordination remains poorly defined. Therefore, we performed Rif-seq experiments in *C. crescentus* to measure genome-wide mRNA lifetimes and analyzed the impact of transcription and translation. Based upon the RNAP elongation speed, we identified that approximately 20% of mRNAs were cotranscriptionally degraded. We generated absolute quantitative estimates of mRNA copy numbers in *C. crescentus*, a useful systems biology resource, revealing that some gene categories have coordinated mRNA turnover. To investigate translation’s impact on mRNA decay, we found that translation efficiency measured by ribosome profiling correlates with mRNA lifetime. We compared the 5’ P cleavage sites to ribosome occupancy and found that cleavage sites occur preferentially in regions of low ribosome occupancy. Using the translation initiation inhibitor retapamulin, which traps ribosomes at the start codon, and chloramphenicol, which arrests elongating ribosomes, shows that chloramphenicol leads to global mRNA stabilization. Surprisingly, we find that the codon adaptation index is inversely correlated with mRNA lifetime, suggesting slow translation elongation may be stabilizing mRNAs from decay. We confirmed the roles of translation initiation and elongation on mRNA lifetimes by generating synthetic YFP and mCherry reporter mRNAs. Taken together, mRNA decay is globally interconnected with transcription and translation.

**Highlights:**

● Quantitative analysis yields absolute mRNA abundance and half-lives for *C. crescentus*
● Identification of 47 new stable ncRNAs
● Both transcription and translation are globally coordinated with mRNA decay
● Translation initiation and elongation impact mRNA decay through ribosome occupancy

## Introduction

Gene expression is an essential process, and in bacterial cells is spatially coordinated within the cytoplasm. While mRNA transcription and mRNA translation have been studied extensively, less is known about the coordinated global regulation of mRNA decay. Most studies of mRNA decay in bacteria have been performed in *E. coli*, however, the localization pattern of mRNA decay machinery is distributed with different localization patterns in different species^1,2^.

The major mRNA decay enzyme, RNase E is the most common scaffold for the RNA degradosome^3,4^. RNase E is localized to the inner membrane in *E. coli*, where it is localized away from the nucleoid where RNAP transcribes mRNA^5^. Co-transcriptional mRNA degradation is suggested to be very rare due to the spatial separation^6^. Conversely the RNase E of other species of bacteria is localized in the cytoplasm, such as *C. crescentus*^1^. While *lacZ* mRNA was shown to be co-transcriptionally degraded in *C. crescentus*^6^, the potential for coordination genome-wide has not been fully explored in this organism. In addition, while a large number of stable non-coding RNAs have been identified in various bacteria^7,8^, Rif-seq has the potential to allow for mapping of stable ncRNAs across the transcriptome, yet Rif-seq has not been directly applied to mapping stable ncRNAs.

To build a systems-wide map of bacterial gene expression, an important goal is to know the absolute copy number and biosynthesis rates for cellular mRNAs. Initial estimates of bacterial mRNA copy number distributions revealed that most mRNAs are present at a copy number of <1/cell, indicating that there is potentially cell-to-cell heterogeneity in mRNA expression^9,10^. While absolute mRNA copy numbers can aid efforts to perform systems-wide modelling of bacterial gene expression networks, absolute biosynthesis and decay rates can further help to build system wide models of bacterial gene expression programs^11^. Despite its use as a model for bacterial systems biology^12^, absolute copy number, transcription rate, and mRNA half-life data is not broadly available in *C. crescentus*.

Finally, mRNA translation is known to modulate mRNA lifetimes across various organisms, suggesting these processes are strongly coordinated. In yeast, translation elongation has been found to have a strong impact on mRNA decay, where optimal codons have been found to protect mRNAs from decay^13^. Additionally, estimates of translation initiation rate have also been found to globally impact mRNA decay^14^, suggesting that this step in translation is also critical in impacting mRNA decay. In *E. coli*, recombinant protein expression libraries showed that codon usage was correlated with mRNA abundance and lifetime^15^, suggesting that translation elongation may be implicated in protecting mRNA like yeast. However, unlike *E. coli*, the *C. crescentus* transcriptome is dominated by non-SD initiated ribosome binding sites^16^, suggesting it predominantly uses different modes of translation initiation than *E. coli*. While *C. crescentus* was one of the first bacteria with ribosome profiling data^17^, no systematic studies have investigated the global relationship between translation and mRNA decay in *C. crescentus*.

To explore the global coordination with transcription and translation with mRNA decay in *C. crescentus*, we performed Rif-seq in *C. crescentus*. By comparing the transcription elongation rate with the mRNA lifetimes, we observed 20% fraction of the transcriptome undergoes co-transcriptional mRNA decay, likely aided by the cytoplasmic localization of *C. crescentus* RNase E^1^. An important aspect of rifampicin induced transcriptional shutoff is that unstable RNAs are rapidly degraded, allowing the mapping of stable RNAs. Therefore, we also utilized this approach to map novel stable RNAs expressed from the *C. crescentus* genome. By measuring the absolute mRNA copy number, we calculated the absolute mRNA transcription rate (molecules/cell/minute) and found that some functional gene categories have distinct mRNA turnover rates, with ribosomal protein mRNAs being rapidly turned over, while other categories like flagellin mRNAs were turned over more slowly. The absolute RNA copy number, mRNA half-lives, and mRNA transcription rates will provide a useful resource for systems biology modelling of the *C. crescentus* cell cycle and asymmetric cell division.

To investigate the relationship between mRNA translation on mRNA decay, we compared ribosome profiling data to mRNA half-lives and found that translation efficiency is correlated with mRNA half-life. Using 5’ P end sequencing to identify RNA cleavage sites, we found that mRNA cleavage sites predominantly occur in regions of low ribosome occupancy, suggesting that ribosomes may directly protect mRNAs from endonuclease cleavage. To probe the roles of translation initiation and elongation on mRNA half-life, we performed Rif-seq in retapamulin (initiation inhibitor) and chloramphenicol (elongation inhibitor) treated cells, revealing that while retapamulin led to a slight increase in mRNA half-lives, chloramphenicol dramatically stabilized the entire transcriptome, suggesting ribosome occupancy along the CDS is strongly stabilizing. By comparing the codon adaptation index, a measure of elongation speed, to mRNA half-life, we saw an inverse correlation, suggesting that slow-moving ribosomes may be protecting mRNAs in *C. crescentus* (in contrast with prior data in yeast^13^ and *E. coli*^15^). To explore the roles of translation initiation and elongation on mRNA decay in a systematic manner, we synthesized synthetic YFP and mCherry genes with various CAI values across the natural range in combination with either a strong or weak translation initiation site. We observed that low CAI and strong translation initiation sites led to mRNA stabilization, while high CAI and weak translation initiation site led to mRNA destabilization, suggesting that both translation initiation and elongation contribute to mRNA lifetime.

## Results

### Global measurement of mRNA half-life

To measure mRNA half-lives across the *C. crescentus* transcriptome we performed Rif-seq, which stops transcription initiation with the antibiotic rifampicin and allows measurement of mRNA lifetime by RNA-seq at specific timepoints after transcription inhibition (Fig 1A). Samples were taken at 1, 2, 4, 8, and 15 minutes after rifampicin addition in mid-log *C. crescentus* cells grown in M2G minimal media. Importantly, as rifampicin only blocks transcription initiation and not elongation, a lag in mRNA decay can often be observed in longer mRNAs (Fig 1B). To account for this lag, which is a result of RNAP runoff, we used the Rifcorrect package^18^ which calculates the runoff time based on the average RNAP elongation rate and distance of a given CDS from the TSS. When examining the Rifcorrect curve fits of our replicate Rif-seq datasets, we found a strong correlation between the half-life values of the two biological replicates (R^2^=0.81, Fig 1C). However, as many mRNAs can have alternative mRNA isoforms, arising from processing sites or alternative TSSs or terminators, which could lead to differences in half-lives, we focused on “simple” mRNAs, which are known to contain only a single CDS and lack alternative TSSs, alternative terminators, or processing sites^19^ (Fig 1C). Importantly, an up-to-date annotation of the transcriptional units in the *C. crescentus* genome was available to allow for the differentiation of complex and simple operons^19^. Upon examining, the correlation of mRNA half-life values was improved when restricting analysis to only simple mRNAs (R^2^=0.84) (Fig 1C), suggesting rather reliable half-life measurement by these datasets. In total using Rif-seq, we were able to measure the mRNA half-lives of 1439 mRNAs across the 3885 annotated CDSs in the *C. crescentus* genome, while only a few mRNAs had half-lives longer than the Rif-seq timepoints used, preventing half-life determination, the vast majority of mRNAs that were not quantified were low abundance transcripts with half-lives that were too fast to be measured by the RNA extraction timepoints in our dataset.

**Figure 1:**
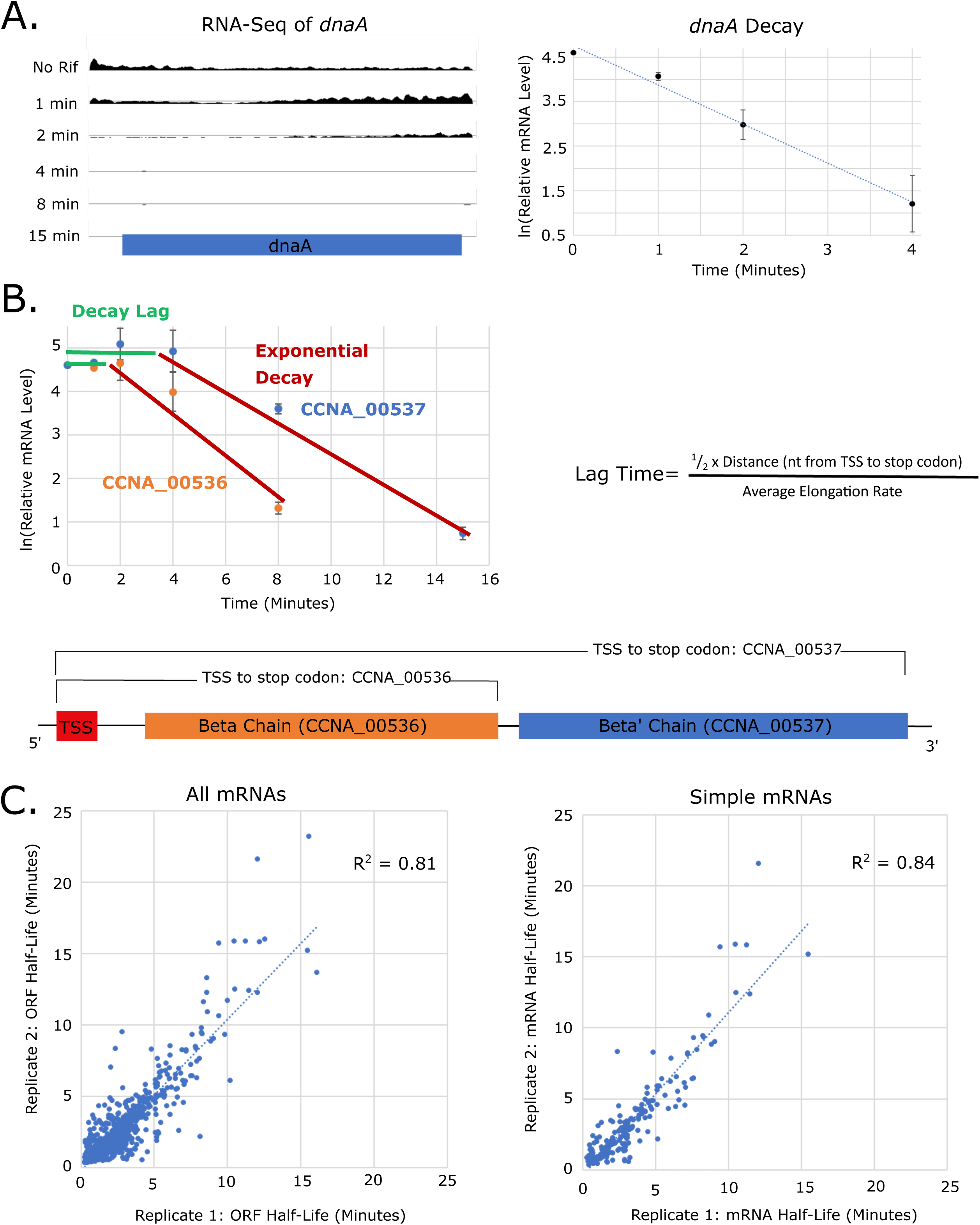
Global quantitation of *C. crescentus* mRNA half-lives by Rif-Seq. A) <u>Rif-seq representative data for a simple short mRNA.</u> (left) Raw read data along the length of the mRNA for *dnaA* gene for each time point. (Right) Log transformed RPKM data plotted against time for the *dnaA* gene. The slope from the linear regression of the log transformed RPKM data was used to find the half-life. B) <u>Rif-seq representative data for a</u> <u>long polycistronic mRNA is impacted by RNAP runoff.</u> Rifampicin inhibits polymerase transcription initiation, but not elongation. This introduces delay before long mRNAs decay as shown with the CCNA_00536/CCNA_00537 operon. Decay lag was corrected for by calculating the average time it takes for the polymerase to run off a particular gene, calculated as the lag time using Rif-correct package^18^. C) <u>Rif-seq produces reproducible mRNA half-lives across biological</u> <u>replicate datasets.</u> (left) For all mRNAs measured in both Rif-Seq biological replicates the half-lives were plotted against each other. (right) For simple (monocistronic and single mRNA isoform) genes found in both Rif-Seq biological replicates the half-lives were plotted against each other. There was a better correlation between replicates with simple genes than with all genes.

### Identification of stable RNAs

Upon examining the mRNAs whose half-lives varied by more than 50% between replicate datasets, we noticed that some of these RNAs contained unannotated ncRNAs that appeared to be processed out of the mRNAs and were stable after 15 minutes after transcription shutoff. We therefore examined the Rif-seq dataset to identify stable RNAs across the transcriptome. To test whether the Rif-seq data could identify stable processed ncRNAs, we examined the abundant tRNAs, as tRNAs are known to be processed at both the 5’ and 3’ ends and easily identified (Fig 2A). By searching for continuous stretches of read density above 15 reads/nt, we found 40 out of 40 tRNAs could be identified with nt level accuracy in the 15’ post rifampicin dataset, suggesting that this dataset can indeed be used to identify stable ncRNAs. We then used this method to search the rest of the *C. crescentus* genome for stable RNAs arising from anywhere in the genome. In total, we identified 147 ncRNAs, including 45 new stable ncRNAs that were not previously annotated (Fig 2B). The new ncRNAs appear to be processed from different parts of the mRNAs (5’ UTR, CDS, or 3’ UTR), transcribed from intergenic regions, or present antisense to another gene (Fig 2B,C). Those mRNAs who overlap with stable ncRNAs were categorized in the “complex” category, and not used for further analysis. The identify of these ncRNAs has been included in the latest NA1000 genbank entry, with the “stable ncRNA” flag, and can also be found in our updated NA1000 transcriptome map^19^.

**Figure 2:**
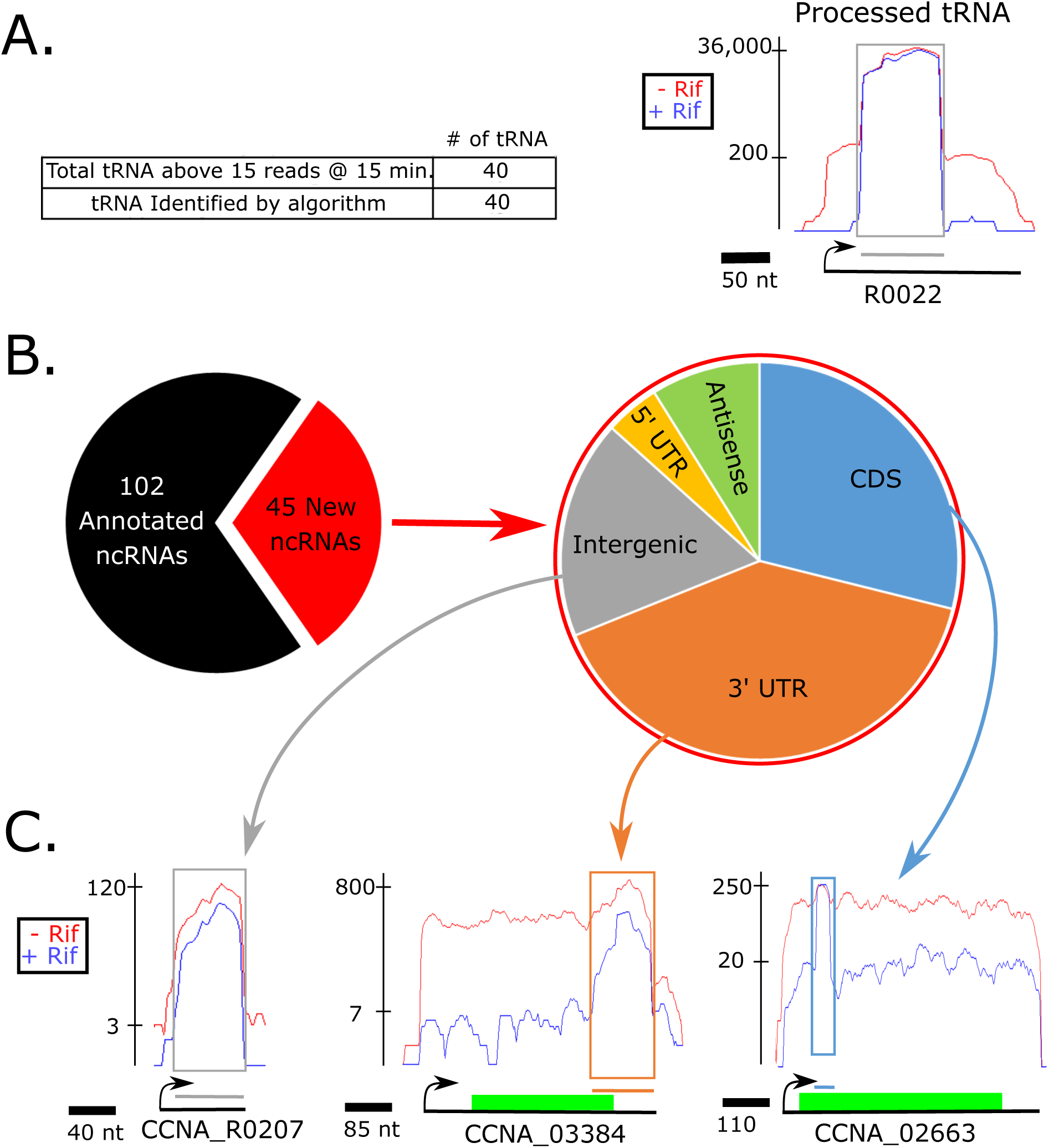
Processed noncoding RNAs are generated from multiple sources. A) <u>Rif-seq can identify stable non-coding RNAs.</u> An algorithm was designed to detect stable mRNA fragments that remain 15 minutes after transcription is inhibited with rifampicin. This algorithm was tested on tRNA and detected all of the processed tRNAs present in the dataset. B) <u>Rif-seq identified 45 new stable non-coding RNAs.</u> There are 102 previously annotated ncRNA in *C. crescentus*, and we detected 45 new stable ncRNA. These new ncRNA were found in a variety of sources across the genome. C) <u>Representative stable non-coding RNAs arising from CDS, 3’ UTR, or intergenic regions.</u> CCNA_R0207 is an example of a new intergenic ncRNA. CCNA_03384 has an example of a new ncRNA that comes is derived predominantly from a 3’ UTR. CCNA_02663 is a new ncRNA that is processed from a CDS. Green bars represent the annotated CDS region, while grey bars represents the newly identified intergenic ncRNA.

### mRNA decay is coordinated with transcription

Since rifampicin only blocks transcription initiation and not elongation, we examined the rates of RNAP runoff of long mRNAs. Indeed, by exploring mRNA transcriptional units above 7kb, we find that the average RNAP density drops at the 5’ end corresponding to the time of transcription initiation shutoff (Fig 3A). By extracting the ½ maximal median read density at each time point after transcription initiation shutoff, we used a linear regression of the distance that RNAP traveled over time, yielding an estimated RNAP elongation rate of 19.37 nt/sec averaged across long operon mRNAs (Fig 3B). This *C. crescentus* RNAP elongation rate is slower than the *E. coli* RNAP elongation rate for mRNAs when grown in LB at 37°C, (42nt/sec^20^), but close to the elongation rates measured in LB at 30°C (25 nt/s^21^) or M9 glycerol at 30°C (20-30nt/s^22^). Using the average RNAP elongation rate, we calculated the transcription times for all the simple mRNAs across the *C. crescentus* genome and compared them with their lifetimes (Fig 3C, Table S1). Here, we found a significant fraction of mRNAs have a lifetime that is shorter than the calculated elongation time, suggesting they are degraded cotranscriptionally (73/375). While evidence for co-transcriptional mRNA decay in *C. crescentus* was reported for a single gene, *lacZ*^6^, our Rif-seq shows that this mechanism is broadly utilized across the transcriptome, likely aided by the cytoplasmic localization of the *C. crescentus*.

**Figure 3:**
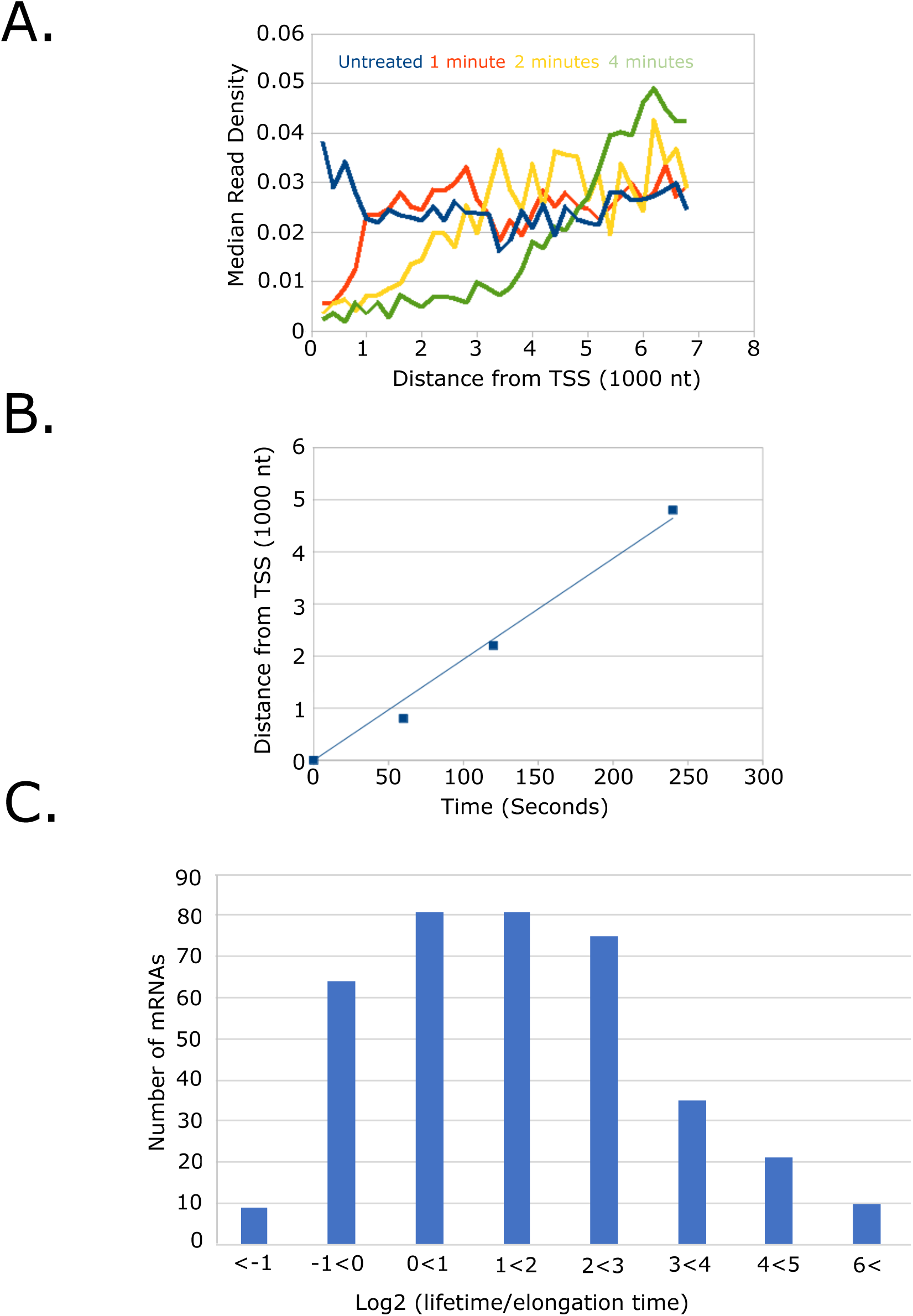
Co-transcriptional mRNA decay occurs in a significant fraction of mRNAs. A) Global runoff of RNAP after rifampicin treatment. Combined read data for highly expressed operons longer than 7,000 nt, at 0, 1, 2, and 4 minutes after the addition of Rifampicin was plotted as distance from the TSS in 300 nt bins versus median read density. **B)** <u>Calculation of RNAP speed from global runoff.</u> The nucleotide distance from the TSS with half-max median read-density was calculated at each time point from the data in panel A, indicative of the average RNAP distance traveled at that time point. The time was then plotted against the calculated half-maximum distance, for each time point. A linear regression was performed on these data and the slope of this regression represents the average elongation rate (19.37 nt/s). **C)** <u>Global estimation of co-transcriptional mRNA decay.</u> mRNAs with a Log_2_(lifetime/ elongation time) less than 0 have lifetimes that are shorter than their average elongation time indicating that these genes should undergo co-transcriptional degradation. While a majority of mRNAs have lifetimes longer than their transcription time, a significant portion of mRNAs have a shorter lifetime, indicative of co-transcriptional degradation.

To better understand the relative importance of transcription and mRNA decay in determining mRNA levels, we performed absolute quantitation of *C. crescentus* mRNA abundances. As RNA-seq provides a genome-wide relative measure of mRNA abundance, we first determined the absolute mRNA copy numbers of two *C. crescentus* mRNAs using single molecule mRNA FISH (Fig 4A). Here we utilized 12 Cy5-labeled probes tiling YFP sequence and a xylose-inducible YFP construct (JS86) for single-molecule control, where we can obtain FISH intensity coming from one yfp transcript under a repressed condition. We collected data for one high abundance mRNA (*Hu-YFP, 2.57 copies/cell +-0.04 (ste)*) and one medium abundance mRNA (*cckA-YFP, 0.252 copies/cell +-0.009*) (Fig 4A). Using these measurements, we performed a linear regression on these values compared to the RPKM values from the RNA-seq data and generated a conversion to the absolute mRNA copy number. The absolute mRNA copy numbers calculated from the smFISH calibration curve were rather close to estimates of the mRNA copy number predicted by assuming an average of 1400 mRNA per cell as in ^10,23^ (Fig S1). Across the transcriptome, we find the highest abundance mRNA (*rsaA*) being present at 25 copies per cell (Fig 4B). While RsaA is the major protein making up the *C. crescentus* S-layer, which is needed in very high abundance^24^, all other mRNAs were present at lower copy numbers. The distribution of mRNA copy numbers was quite broad, with the average copy number of 0.087 copies/cell, and a standard deviation of 0.269. Only a small fraction of mRNAs, 40 total, were not detected at all in this growth condition, suggesting that most genes in the *C. crescentus* chromosome are expressed during logarithmic growth.

**Figure 4:**
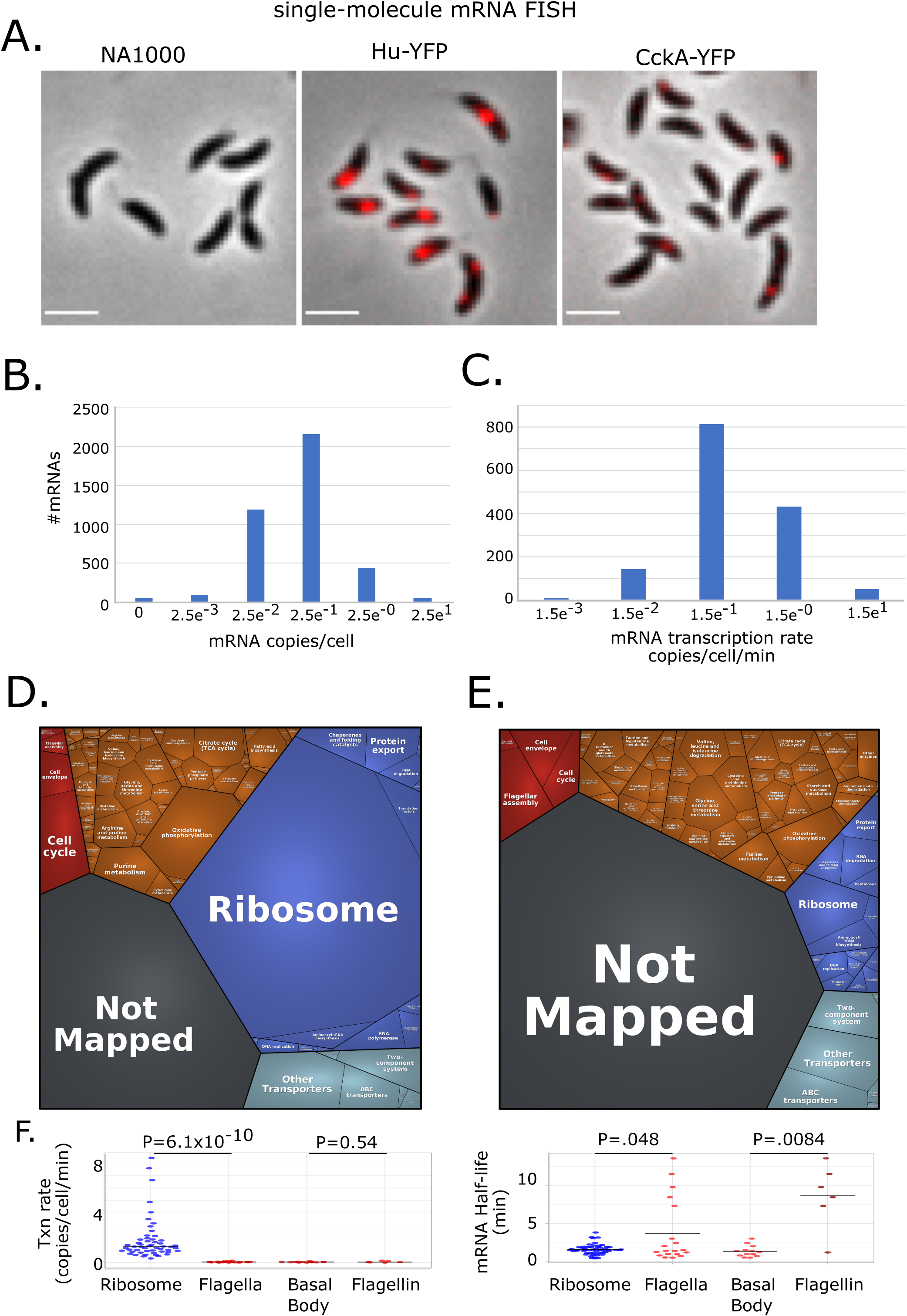
Steady state mRNA levels are determined by a combination of transcription and decay. A) <u>Absolute mRNA levels were determined by single-molecule mRNA FISH using anti-YFP FISH probes.</u> The mRNA abundance of the Hu-YFP mRNA (*2.57 copies/cell*) and CckA-YFP mRNA (*0.252 copies/cell*) were determined. B) <u>Distribution of absolute mRNA levels of the *C. crescentus* transcriptome.</u> Absolute mRNA levels were calculated from the relative abundance measurement by RNA-seq via the measured absolute mRNA abundance values of Hu-YFP and CckA-YFP mRNAs. C) <u>Distribution of absolute mRNA production rates of the *C. crescentus* transcriptome.</u> Absolute mRNA transcription rate in molecules/cell/minute were calculated by dividing the mRNA half-lives by the absolute mRNA copy number. D) Resource allocation of mRNA transcription. Proteomap representing each gene category with area scaled to the calculated mRNA transcription rate in molecules per cell per minute. E) Resource allocation of mRNA decay. Proteomap representing each gene category with area scaled to the calculated mRNA half-lives in minutes. F) Detailed comparison between the ribosome category and flagellar assembly categories. Left, transcription rate data are plotted for each category, and the flagella category is split into both the basal body and flagellin parts of the machinery. Right, mRNA half-life data are plotted for the same categories. P-values are calculated from 2-tailed T-tests with unequal variance. “Corresponding supplementary table:” Example genes CCNA_00808, and CCNA_01826 showing the different expression strategies for two similarly highly expressed genes.

As the steady state mRNA level is determined by both the transcription rate (molecules/cell/minute) and the half-life (minutes), we can calculate the transcription rate based upon the absolute measurement of copy number and mRNA half-life measurements (transcription rate=absolute mRNA copy number(measured)/lifetime (half-life/ln(2)) (measured)). The absolute transcription rates varied from 8.4 mRNAs/minute, with many ribosomal protein mRNAs among the top transcribed genes, to 0.00055 mRNAs/minute (Fig 4C, Table S2). To visualize the data, we generated a proteomap of the calculated absolute transcription rates, with each polygon’s area representing the summed transcription rates of all the mRNA representing each category (Fig 4D). The area of each polygon represents the load of RNAP toward their transcription. Here we find that indeed, ribosomal protein mRNAs are transcribed at the highest rate, which is consistent with the needs of nascent ribosomes in logarithmically growing bacteria^25^. Other categories of genes, for example, flagellar assembly, have a much lower summed transcription rate. We also generated a proteomap of the mRNA half-lives to allow us to compare the rates of mRNA decay the same classes of mRNAs (Fig 4E). Here we observe the converse, where ribosomal protein mRNAs have quite short mRNA lifetimes, while other classes of mRNAs like flagellar assembly were observed to have much higher mRNA lifetimes (Fig 4F). When examining the rates of transcription for all mRNAs in each category, we indeed observed that all ribosomal protein mRNAs have higher transcription rates as compared to flagellar assembly mRNAs, however, only a subset of flagellar mRNAs had long half-lives (Fig 4F). Importantly, flagellin mRNAs were generally quite stable^26^, which are needed at very high copy number to build the flagellar filament, while the mRNAs encoding basal body proteins were generally less stable (Fig 4F). Overall, this presents two distinct strategies to yield the desired mRNA copy number the cell needs. Ribosomal proteins mRNAs are abundant through high transcription rates, but have short half-lives, presumably to allow rapid shutoff in the event of a sudden changes in growth condition. Conversely, flagellin mRNAs are quite abundant, but are transcribed at a rather low rate and are therefore stable from mRNA degradation, potentially allowing the cell to maintain their flagella even under conditions of nutrient limitation. As this apparatus can be used to swim towards gradients of higher nutrients, the slower turnover of the flagellin mRNAs may be an advantageous adaptation under such nutrient depleted conditions. Therefore the absolute quantitation of mRNA abundance, calculated transcription rates, and half-lives will provide a useful dataset to better understand the role of transcription and mRNA turnover in setting the optimal mRNA levels for bacteria to optimize their fitness in their ever changing environments.

### mRNA decay is globally coordinated with translation

We next investigated the potential global role of coordination of mRNA decay and translation. As mRNA decay in many bacteria initiates with endonucleolytic cleavage by RNase E yielding 5’ P sites^4,27^, we first investigated the pattern of 5’ P sites present in the *C. crescentus* transcriptome. To identify sites of endonucleolytic cleavage, we utilized a previous 5’ global race dataset in which total RNA or tobacco acid pyrophosphatase (TAP) treated total RNA were subjected to 5’ end sequencing^17,28^. As depicted for a tRNA gene in Fig 5A, increased reads in the Tap treated library is indicative of the TSS, while reads that are depleted in the 5’ P sites are indicative of endonuclease cut sites, in this case by RNase P which processes the 5’ end of tRNAs. Indeed, the log_2_ ratio of reads in TAP (+/-) samples can be used to distinguish 5’ PPP sites from 5’ P sites (Fig 5A). As prior analysis only focused on 5’ PPP sites^28^, here we used the 5’ P sites across the *C. crescentus* transcriptome. Across all 19,356 5’ P sites identified, we observed a cut site motif with a slight preference for an A at the cut site, and a preference for a U 1 nt away from the 5’ end (Fig 5B), which is very similar to the RNase E cut site motif determined by cross-linking RNAs directly to RNase E in *Salmonella*^29^. As the global RACE method does not enrich transcripts for RNase E cleavage site, this suggests that RNase E likely is the dominant mRNA decay nuclease generating 5’P ends in *C. crescentus.* Analysis of 5’ P cut sites also revealed that these sites contain a lower average predicted secondary structure content (Fig 5C), suggesting cutting occurs in regions of the mRNAs that are more likely to be single stranded. We then analyzed the locations of the mRNA cut sites within mRNAs (Fig 5D). Here we find most cut sites (∼4000) are present in the CDS, followed by the 5’ UTR (∼1100), near the start codon (∼500), near the stop codon (∼250), and in the 3’ UTR (∼50). As a reference for we also examined the cut site density of 5’ P sites tRNAs, as these RNA’s are processed by RNaseP to yield mature 5’P ends. As each of these regions vary significantly in size, we also examined the density of cut sites/nt within each of these features. tRNAs which ideally contain one 5’ P site, had the highest 5’ P cutting density, largely due to their relatively small size. 5’ UTRs and around the start codon contain a similar high 5’ P site density, which may be due to a scanning by RNase E from the 5’ end^30^. Despite having the most overall cut sites identified, CDSs conversely show a rather low cut site density (∼4 every 1000nt), which is in part due to the much large size of CDSs relative to UTRs, start codon region, or stop codon region.

**Figure 5:**
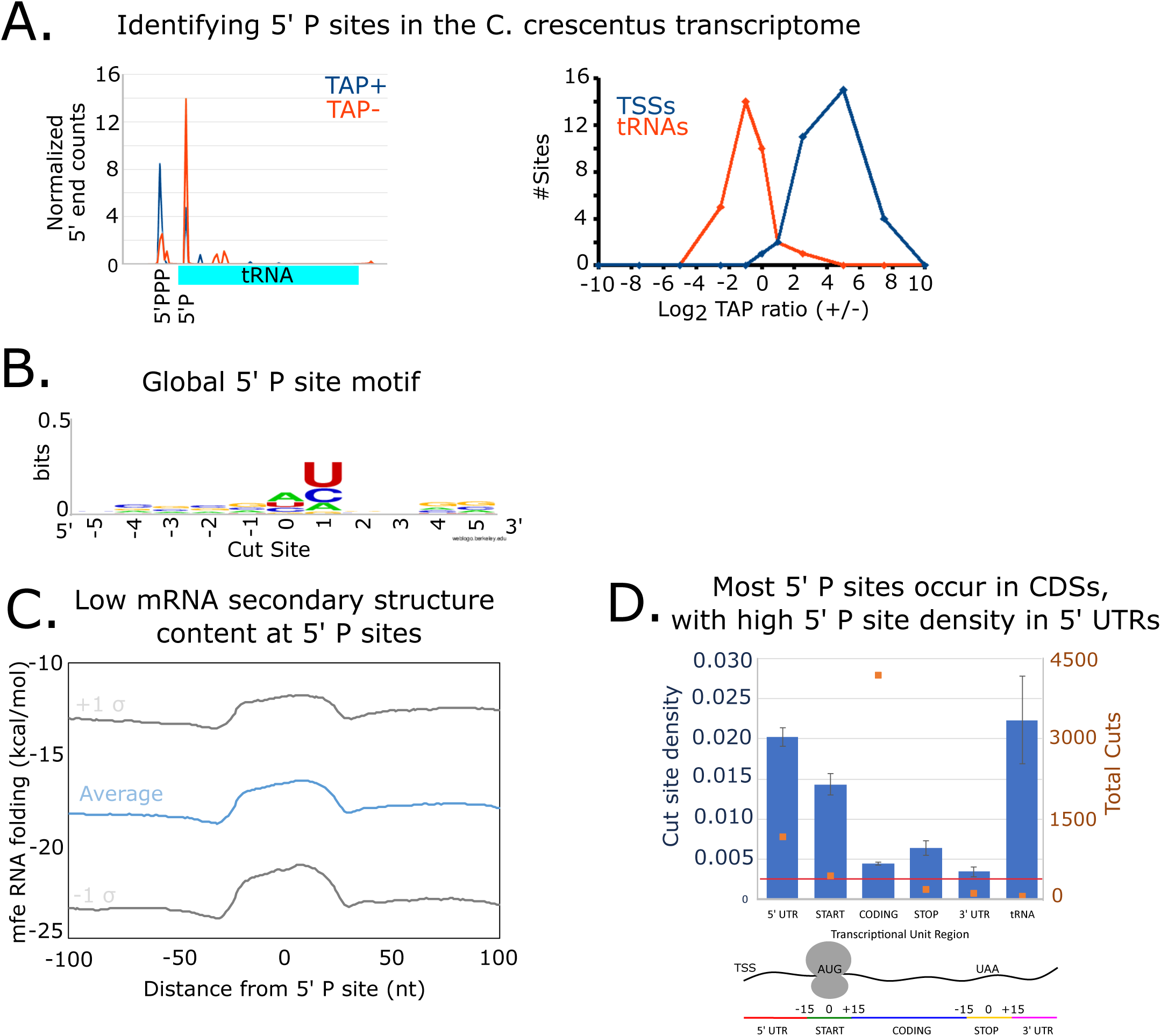
mRNA cut sites are enriched in sites of low structure content in the CDS. A) <u>5’ Global RACE can identify RNA cleavage sites.</u> (Left) 5’ PPP data generated from Tobacco Acid Pyrophosphate (TAP) enriched 5’ global RACE^28^ on tRNA for the TAP + and TAP – negative libraries showing tri-phosphate native ends, and mono-phosphate processed ends. (Right) Log2 Ratio of TAP + library to TAP – library show that processed ends of tRNA have a negative Log2 ratio and primary TSS have a positive Log2 ratio for tRNAs. B) <u>Global motif of RNA cleavage sites.</u> Sequence motif of degradation cut sites from TAP shows a preference for uracil one nucleotide after the cut. This is a similar preference that has been found in other experiments that specifically enriched for RNase E cutting [Fig S3B ^29^]. C) <u>mRNA cleavage sites have reduced secondary structure.</u> Metagene plot of Minimum free energy (mfe) of RNA secondary structure centered on the cut sites. mfe was calculated for each nucleotide around the cut sites with a 50 nucleotide sliding window use the RNAFold package^52^. Standard deviation above and below the average mfe are plotted. D) <u>Global profile of mRNA cleavage sites.</u> On the left y-axis cut site density (cuts/nt) is shown for different regions for mRNA, and for tRNA. On the right y-axis the total cuts per region is shown. There are more total cuts occurring in the CDS region of mRNA, but there is a higher probability of a cut per nucleotide in the 5’ UTR. The red line shows the cut density level if they were randomly distributed across all regions of mRNAs. Error bars are estimated as wald confidence intervals.

To better understand how the patterns of mRNA cut sites might influence translation or vice versa, we next compared ribosome profiling datasets measuring translation efficiency to the mRNA half-lives generated by Rif-seq in the same growth conditions^17^. Here, as observed in other studies ^14,15,31^, there is a positive correlation between the translation efficiency of mRNAs (here it is defined as the ribosome profiling read density divided by the RNA-seq read density) and mRNA half-lives (R^2^ = 0.58). In total, we observe a 4-fold difference in mRNA lifetimes across the different bins of translation efficiency (Fig 6A). To test whether translation initiation or elongation has a larger effect on mRNA lifetimes, we performed Rif-seq experiments with cells treated with the initiation inhibitor retapamulin or the elongation inhibitor chloramphenicol (Fig 6B, Table S4). After treating cells with the initiation inhibitor retapamulin, which is known to trap ribosomes over the start codon, and allow runoff of elongating ribosomes, we observed a slight increase in mRNA half-lives across the transcriptome. We next compared the sites of RNA cleavage with the average position of ribosomes within the mRNA CDS as determined by ribosome profiling (Fig 6C). Because rates of initiation and termination differ from elongation, we removed start and stop codon regions from the analysis. We hypothesized that if ribosomes are able to protect the mRNA from cleavage, we would expect a lower density of ribosomes at cut sites. In line with this hypothesis, we observe that on average, there is an approximately 20% reduction in ribosome density at the RNA cut site (Fig 6C). In line with this, if we examine mRNA CDS sites by their ribosome occupancy, we observe that there is an approximately 2-fold higher probability of finding an RNA cut site at a region in the mRNA of low ribosome density as compared to high ribosome density (Fig. 6D). Direct protection by ribosomes is unlikely to be the only mechanism affecting mRNA lifetime. As shown in figure 6C, we also observed a slight increase in ribosome density directly after the cut site. Indeed, in the heatmap representation below, a small fraction of cut sites appear to have high ribosome density directly after the cut site, which may suggest that some cuts occur in the mRNA at the 5’ side of high occupancy ribosomes.

**Figure 6:**
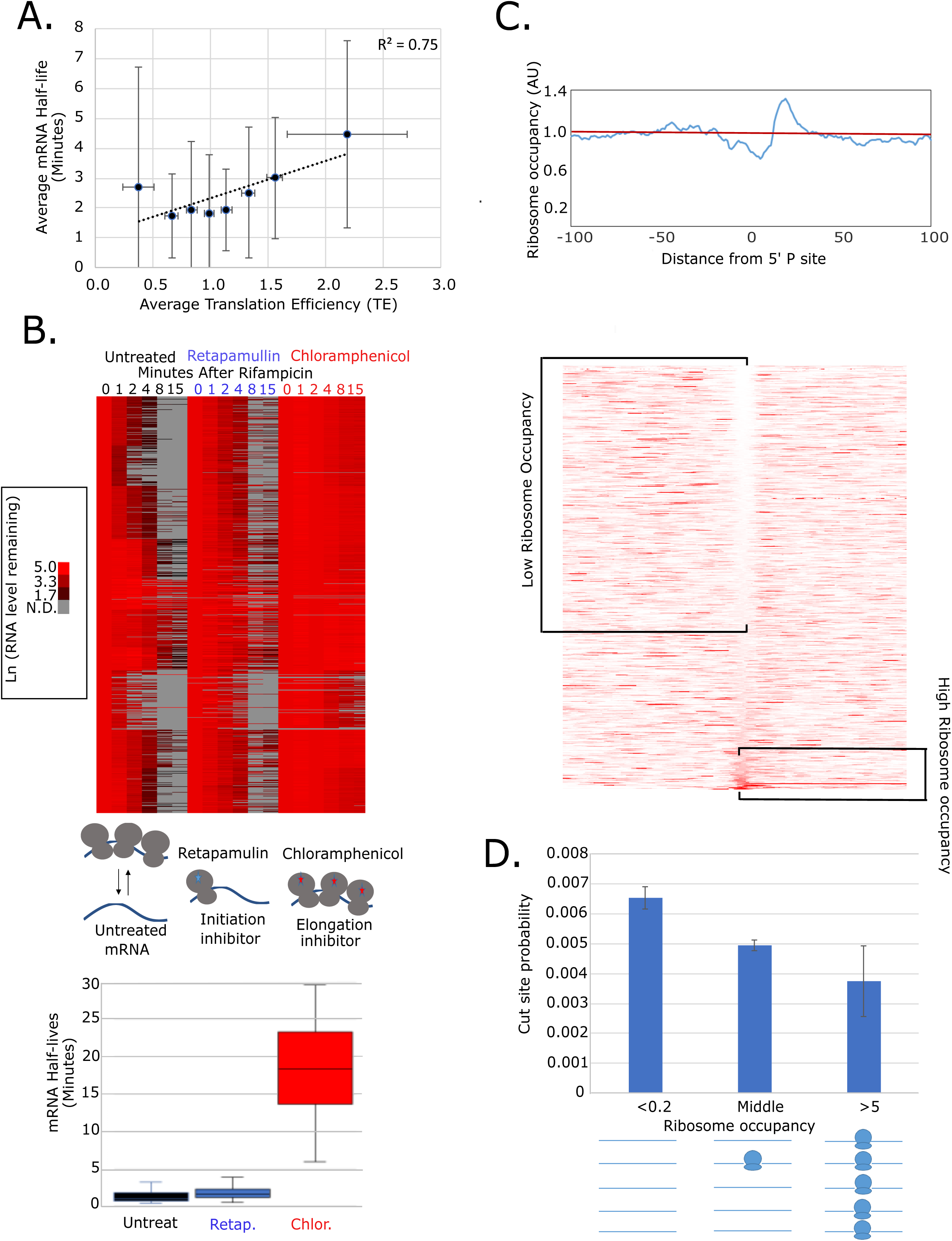
Ribosome protection globally impacts mRNA decay. A) <u>Positive correlation between translation efficiency^17^ and mRNA half-life</u>. mRNA was binned according to their average translation efficiency, and average mRNA half-life. The error bars represent the standard deviation of the mRNA half-lives. A linear regression trend line is shown with its R^2^ value of 0.58. B) <u>Ribosome occupancy is a major determinant of</u> <u>mRNA half-life.</u> (top) mRNA half-lives in minutes were measured using Rif-seq from cells lacking drug treatment, cells pre-treated with, or cells pre-treated with chloramphenicol. Each lines represents a single mRNA with its abundance over time plotted as a heat map with the indicated abundance scale on the left. (bottom) Box-plots of the mRNA half-lives of mRNAs whose half-lives were measured in all three data sets. C) In the CDS, <u>mRNA cleavage typically occurs at</u> <u>sites of low ribosome occupancy.</u> (Top) Meta-gene plot of Distance from cut site is plotted versus ribosome occupancy with the red line showing the average occupancy across all CDSs. (bottom) Ribosome occupancy of each cut site region as a red heat map (more red is higher ribosome occupancy). Each line represents the ribosome occupancy in a 200nt window centered on the 5’P site. D) <u>Cut site probability is inversely correlated with ribosome occupancy.</u> Each nucleotide in CDSs were sorted into 3 categories based on their ribosome occupancy; Cut sites that had ribosome occupancy at least five times less than average (<0.2), cut sites that had ribosome occupancy at least five times more than average (>5), and cut sites in-between. The total cut sites were then summed for each category, and the cut site probability was calculated as number of cut sites/total number of sites. Error bars represent 95% Wald confidence interval.

To explore whether the trends in *C. crescentus* mRNA lifetime are caused by translation elongation or initiation, we first examined the codon adaptation index (CAI) across *C. crescentus* mRNAs (Fig 7A). The CAI is related to the relative abundance of a particular codon across the transcriptome, with a strong correlation of CAI and tRNA abundance, leading to faster translation of codons with high CAI, and slower translation of codons with low CAI. While it would be useful to know the initiation rate of each mRNA, there are currently no algorithms which can accurately calculate the rate of translation initiation. Across *C. crescentus* mRNAs, the CAI values range from 0.34 to 0.89. To examine how CAI relates to translation, we observed a mild positive correlation between CAI and TE with an R^2^ of 0.20 (Fig 7A). Interestingly, we observed a negative correlation between CAI and mRNA half-life (Fig 7A) with a stronger R^2^ of 0.66. The lowest CAI bin appears to lead to a strong stabilization (average half-life = 6.9 min), while the other variants have average half-lives between 2 to 3 minutes. This was surprising, as optimal codons in eukaryotic mRNAs are typically found to stabilize the mRNAs^13^. However, some studies in this system have also implicated translation initiation as exerting major control of mRNA lifetimes^14^.

**Figure 7:**
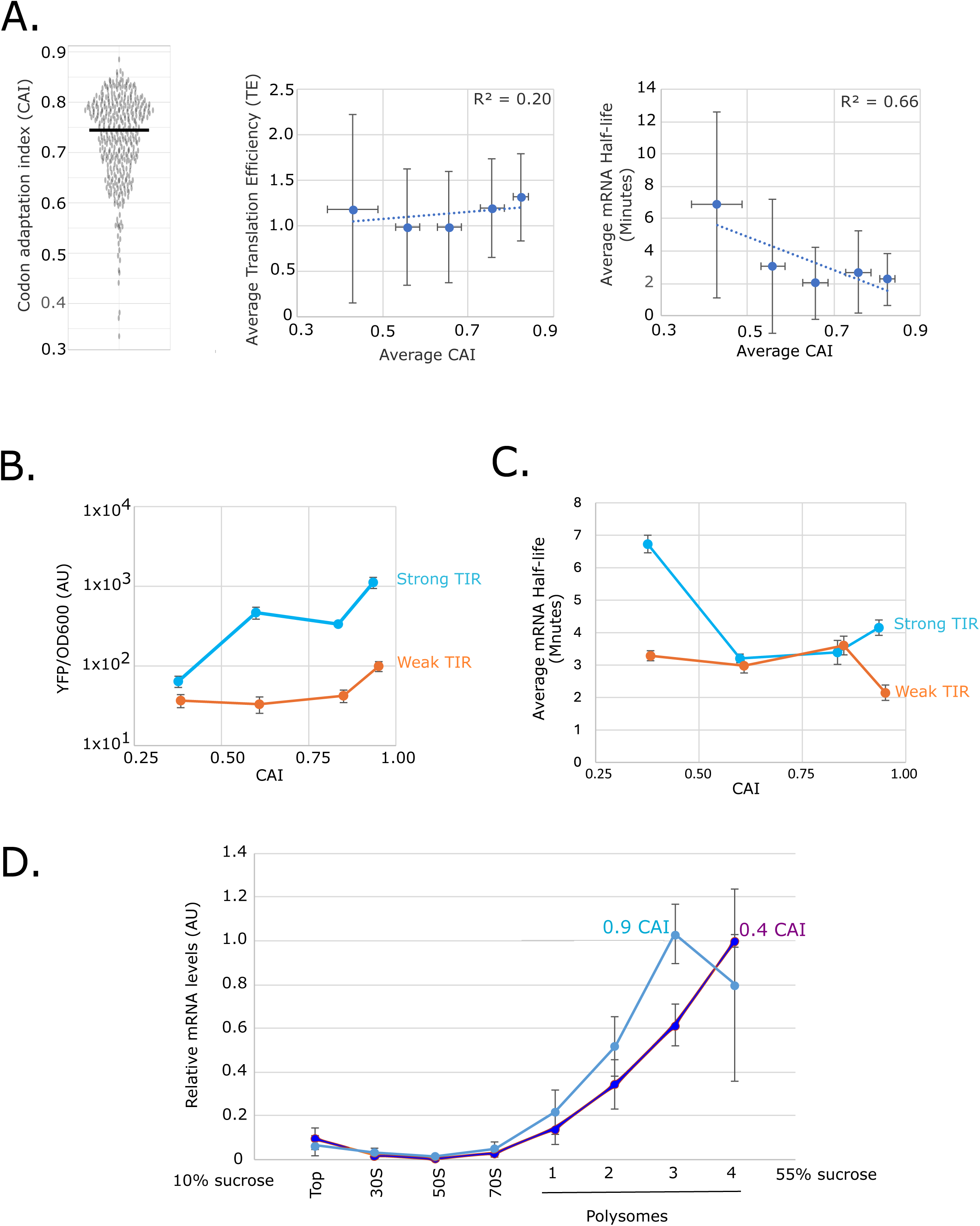
Translation initiation and elongation impact mRNA half-life. A) Codon usage is positively correlated with translation and negatively correlated with mRNA half-life. Left: Codon adaptation index (CAI) for each *C. crescentus* CDS. Middle: simple mRNAs were binned by their CAI values and the translation efficiency^17^ was compared. R^2^ determined by a linear curve fit. Right: simple mRNAs were binned by their CAI values and the mRNA half-life was compared. R^2^ determined by a linear curve fit. Error bars represent standard deviation. B) <u>YFP translation is a result of both initiation and elongation.</u> YFP was measured in a fluorescent plate reader for variants with different translation initiation regions and codon usage frequency (CAI). The YFP intensity (AU) was divided by the optical density as a measure of the YFP/cell. C) <u>Translation initiation and elongation coordinately impact</u> <u>YFP mRNA half-life.</u> mRNA half-life was measured by qRT-PCR for each YFP variant with different translation initiation regions and codon usage frequency (CAI). D) <u>Highest polysome occupancy occurs when strong initiation and poor</u> <u>elongation are combined.</u> 10-55% sucrose gradients were used to separate polysome fractions, and mRNA occupancy across the fractions were measured by qRT-PCR. Average mRNA level in each fraction and standard error are plotted. qRT-PCR levels were normalized for each mRNA based upon the highest occupancy fraction. n=3 for 0.4 CAI, n=2 for 0.9 CAI.

To test the role of CAI and the translation initiation region (TIR) on mRNA decay, we designed YFP mRNAs with various TIRs and codon usage frequencies to systematically alter the rates of translation initiation and elongation. To control the rate of transcription, all the constructs were driven by the xylose promoter. We selected one weak and one strong TIR^16^ and combined these with codon usage ranging from highly suboptimal (CAI = 0.38) to highly-optimal (CAI = 0.94) for the range of *C. crescentus* mRNAs. Across these engineered YFP mRNAs, we observed that the YFP fluorescence varied from as high as 1100 AU, to as low as 36 AU (Fig 7B). We observed that increasing the TIR and CAI both positively contributing to the amount of YFP produced (Fig 7B). CAI had a stronger impact on the variant with a strong TIR, varying over a 17 fold range from the lowest to highest CAI constructs, while it had a smaller impact on the constructs with the weak TIR varying over a 2.7 fold range from lowest to highest CAI constructs (Fig 7B). This suggests that the CAI may have become rate-limiting with the strong TIR, while initiation may be rate limiting in the weak TIR construct. We then measured the mRNA lifetimes of these YFP mRNAs using qRT-PCR and found that with a strong TIR, the lowest CAI variant led to a significant stabilization of mRNA half-life (6.7 min), compared to the 3.2-4.2 min half-lives observed for the higher CAI variants (Fig 7C). This suggests that poor CAI may be causative for the observed slower mRNA lifetimes. Interestingly, with the weak TIR we observed no stabilization with low CAI, and the half-lives across the weak TIR variant remained fast (2.1 – 3.6 min). To confirm the strong TIR low CAI stabilization results were not a result of YFP, we also performed the same experiments in mCherry, and found similar results (Fig S4). This suggests that higher translation initiation rate combined with lower CAI likely yields higher polysome occupancy, which protects mRNAs from RNase cleavage yielding longer mRNA lifetimes. To test this, we performed 10-55% sucrose density gradients to separate polysome fractions, and measured the occupancy of the high TIR 0.4 and 0.9 CAI variants across the different polysome fractions (Fig 7D). We observed that the 0.4 CAI variant peaked in a higher density fraction as compared to the 0.9 CAI variant, suggesting that high TIR combined with low CAI leads to higher polysome occupancy.

## Discussion

### Global resource of *C. crescentus* mRNA stability

Genome-wide absolute mRNA quantitation (Fig 4), together with the absolute synthesis and decay rates of those mRNAs (Figs 1,4), provides a useful resource for modeling gene expression in *C. crescentus*. As *C. crescentus* is a premier model of bacterial systems biology^12^, these data will help to aid in future modeling efforts of its gene expression programs. Together with absolute quantitation of mRNA translation data by ribosome profiling^32^, absolute quantitation of the steps of gene expression are now available from transcription to translation, leaving absolute quantitation of proteolysis rates the only remaining global dataset without global absolute quantitation rates. While regulated proteolysis is a key step in the gene expression programs of *C. crescentus*^33^, however, early proteomics data suggests that a majority of proteins are more stable than the duration of the cell cycle^34^, suggesting protein synthesis provides a larger influence on the levels of most proteins.

By analyzing our Rif-seq data, we identified multiple stable ncRNAs that were not previously annotated. Many stable ncRNAs identified are parts of mRNAs, including presumed processing from 3’ rho-independent terminators, similar to organisms like *Salmonella*^35^, while many appeared to be processed from CDSs and intergenic regions (Figure 2). Rif-seq is a rather simple strategy for mapping stable ncRNAs, as the vast majority of the transcripts are unstable, the remaining sRNAs are very simple to detect as there is very low background of reads outside their boundaries. The functions of the newly identified stable ncRNAs is not yet known, but they may provide important functions to the cell, and have been annotated in our map of the *C. crescentus* transcriptome^19^.

### mRNA decay coordinates with transcription

*C. crescentus* mRNA decay revealed to be coordinated with transcription across the transcriptome, with coordination observed between mRNA synthesis and turnover rates and cotranscriptional degradation (Figs 3, 4). We observed that some gene categories have high rates of transcription and short half-lives, while others have slow rates of transcription but long half-lives (Fig 4). While original observations in yeast mRNA turnover pointed to functional specificity by gene function categories^36^, the roles in *C. crescentus* may suggest that some cellular pathways can be shut off quickly in response to environmental changes, such as ribosomal protein mRNAs. As ribosomal production is one of the most energetically costly pathways in bacteria^23^, this may help to rapidly adapt their expression to conditions of low nutrients or cell stress. Other mRNAs like flagellin mRNAs are more stable, yet they use far less cell resources than ribosome production and help the cells swim toward more favorable environments, suggesting they may provide fitness across various conditions. Therefore, it seems as if the functional specificity of mRNA turnover in bacteria is coordinated with the cell’s program of gene expression to optimize fitness during adaptation to uncertain and changing environmental conditions.

We observed a significant fraction of mRNAs who undergo cotranscriptional decay (Fig 3) suggest a broader coordination between transcription elongation/termination and mRNA decay. These mRNAs undergoing cotranscriptional decay tend to be longer, and likely spend their lifetimes adjacent to the DNA loci they were originated from^1^. Interestingly, in *C. crescentus*, the RNA degradosome proteins are localized largely within the cytoplasm, and are within close proximity to RNAP and the nucleoid^1^, which is likely important feature for cotranscriptional decay. Interestingly, <u>B</u>acterial <u>R</u>ibonucleoprotein <u>bodies</u> (BR-bodies), which are biomolecular condensates that are composed of the RNA degradosome and facilitate mRNA decay^37,38^, were found to contain transcription machinery such as Rho^39^, suggesting coordination between transcription elongation and decay may be occurring through BR-bodies. Cotranscriptional decay is likely to vary between species of bacteria, as *E. coli* appears to have membrane anchored RNA-Degradosomes^40^ and BR-bodies^41^ with coupling between transcription and translation, while *B. subtilis* has membrane anchored RNA-Degradosomes and BR-bodies^42^ but does not couple transcription and translation^43^. In fact, when *lacZ* mRNA decay was examined, its co-transcriptional decay was rare in *E. coli* and *B. subtilis* but occurred in *C. crescentus*^6^. An important goal moving forward is to understand how mRNA transcription and decay are globally coordinated across species, and how the observed differences in RNA degradosomes, transcription and translation machinery, and localization of these factors contribute to coordinated decay.

### mRNA decay coordinates with translation

We observed a global correlation between translation efficiency and mRNA half-life, suggesting translation may be protecting mRNAs from decay. As it has long been known that ribosomes protect a footprint on the mRNA^44^, we observed that via chloramphenicol treatment which increases ribosome occupancy on mRNAs, that there is a transcriptome wide stabilization of mRNAs (Fig 5). Conversely, treating cells with retapamulin, which reduces ribosome occupancy, led to little or no change in mRNA lifetimes (Fig 5). In yeast, treatments with hippuristanol and cycloheximide led to similar effects on mRNA decay^14^, both supporting a model in which mRNA is protected by coverage with ribosomes. In line with the role of ribosome protection, we observed that most 5’ P cut sites identified in the genome tend to occur in the CDS, and more specifically in locations with low structure content and low ribosome occupancy (Fig 5, 6). Interestingly, while it was previously found in other organisms that codon optimality, which promotes faster translation elongation rates, could promote longer mRNA lifetimes^13,15,45^, however, we observed an opposite trend in *C. crescentus*, where CAI was anti-correlated with mRNA lifetime (Fig 7). By use of orthogonal synthetic gene constructs, we found that mRNAs with strong translation initiation sites and low CAI appear to be specifically stabilized by having increased polysome occupancy, while strongly initiated mRNAs with high CAI dramatically increased translation but reduced polysome occupancy and were less stable (Fig 7). Codon optimality driven stabilization in eukaryotic species is thought to be driven by the Ccr4-Not complex^46^, which promotes deadenylation of 3’ poly-A tails by recognition of empty E-site in the ribosome. Interestingly, *C. crescentus* lacks Ccr4-Not and the mRNA decay machinery is thought to engage by endonucleolytic cleavage in the mRNA by RNase E^4,27^, suggesting the different mRNA decay pathways may be responsible for the differences in codon usage between organisms.

### Materials and methods

#### Rif-Seq Cell Harvesting

25 mL of *C. crescentus* cells were grown overnight to mid-log phase in M2G media. Cells were grown in an incubator shaker at 28°C 250 rpm in a 250 mL conical flask. Mid-log phase was confirmed as an optical density OD_600_ reading between 0.3 to 0.5. 2 mL of Qiagen RNAprotect bacteria reagent is brought to room temperature in 15 mL falcon tubes. Rifampicin dissolved in DMSO is added to the 25 mL culture at a final concentration of 200 μg/mL. 1 mL of culture is pulled out of the conical flask at each time point after the addition of rifampicin and added to 2 mL of RNAprotect and vortexed for 5 seconds to stabilize mRNAs. Timepoints of RNA collection were 0, 1, 2, 4, 8, and 15 minutes after rifampicin addition. After the last time point was taken, the cells are incubated at room temperature for at least 5 minutes in the RNAprotect. The cells were then spun down and pelleted in preparation for RNA Extraction by trizol. RNA was extracted according to protocol^38^ and libraries were constructed according to a published protocol^47^. RNA-seq libraries were deposited into NCBI GEO with accession number GSE157432.

#### mRNA Half-life calculation using Rif-Correct Software

Half-life for each gene was calculated from Rif-Seq according to the protocol in (Aretakis & Schrader BioRxiv^18^). To calculate lag time (or RNAP runoff time), RNAP elongation time was calculated to the middle of each CDS feature. This distance was calculated from half the distance from the farthest upstream TSS site, or first CDS start codon if no TSS is known (for operons this would be the first gene in the operon), to the end of the CDS site for that specific gene. This was used as a correction, and the linear regression of the natural log of the RPKM data gives a slope calculation which represents the mRNA half-life.

#### Detection of non-coding RNAs

57 non-coding RNAs were discovered by searching the 15’ post rifampicin samples for contiguous stretches of RNA above 15 reads/nt. New ncRNAs were categorized as CDS if they overlapped with the CDS region, as 3’ UTR or 5’ UTR if they overlapped with a previously known UTR region or overlapped with the region within 30 nucleotides of an upstream or downstream start or stop codon, antisense if they overlapped with the region on the opposite strand of a CDS, or intergenic if they met none of the previous criteria and where not overlapping a known CDS or ncRNA. These stable ncRNAs were reported in our previous work^19^ and have been updated to the current version of the NA1000 genome in genbank (NC_011916.1).

#### Calculating average RNAP elongation rate on mRNAs

For each time point (0, 1, 2, and 4 minutes) read data was combined for highly expressed operons greater than or equal to 7,000 bases in length in 300 nucleotide bins. The half-max distance for each timepoint was calculated. Then each time point in seconds was plotted against the half-max, and a linear regression was performed, and the slope of the linear regression corresponded to the average RNAP elongation rate on mRNA in nucleotides per second (19.37 nt/sec). 33 mRNAs >7kB were used in this analysis.

#### Half-Life vs Elongation Time

Elongation time was only calculated for genes that had non-complex mRNA such as those genes that are monocistronic, and without multiple known isoforms (multiple TSS’s or internal TSS’s). Elongation time was calculated as the length of the gene in nucleotides (from known TSS or start of CDS to end of CDS) divided by the average RNAP elongation rate on mRNA’s (19.37 nt./sec). mRNA lifetimes and the comparison with transcription elongation times of all simple mRNAs across the C. crescentus genome is available in Table S1.

#### smFISH

*C. crescentus* cells were grown in M2G at 28°C until OD660 becomes 0.2, and *E. coli* cells were grown in M9 minimal media supplemented with glucose at 37°C until OD_660_ becomes 0.2. For both species, we followed the same smFISH protocol previously published^48^. In short, 0.9 mL of cell culture was withdrawn and immediately mixed with 0.3 mL of 16% formaldehyde for fixation. Cells were adhered to a poly-L-lysine treated coverslip surface and permeabilized by 70% ethanol. For *yfp* mRNA labeling, we used 12 tiling probes for *yfp*, labeled by a single Cy5 at the 5’ end. The sequences are provided in Fig S6. The probe hybridization solution included 20% formamide, 2X saline-sodium citrate (SSC), and 10 nM of the Cy5-labeled probes. The hybridization was performed for 2 hours at 37°C, followed by washing with a solution of 25% formamide and 2X SSC. For imaging, we used a Nikon Eclipse Ti-2 microscope equipped with a 640 nm laser, a phase-contrast objective Plan Apochromat 100x/1.45 NA (Nikon), and an Andor iXon Ultra 897 EMCCD camera. For analysis, we used MicrobeTracker for cell outlines and spot identification^49^.

We used the wild-type strains (NA1000 and MG1655) as a negative control for *yfp* probes. For a single-molecule control, we constructed an *E. coli* strain with a chromosomal copy of *tsr*-*yfp* under the lac promoter (using the same *yfp* sequence as used in *C. crescentus* strains; SK346). A previous study showed that *tsr-venus* mRNA is present as 0.039 copy per cell (or about 4% cells have 1 mRNA) without IPTG induction when cells were grown at 37°C in M9 glucose. We also used a *C. crescentus* strain expressing *yfp* from a xylose-inducible promoter (JS86) as a single-molecule control. *yfp* FISH showed diffraction-limited spots (foci) with comparable intensities in both *E. coli* and *C. crescentus* controls—with *E. coli* showing 2.3% cells with foci while *C. crescentus* showing 12.6% cells with foci (average mRNA number to be 0.03 ± 0.006 in *E. coli* (SK346) and 0.14 ± 0.007 in *C. crescentus* (JS86); Fig S5). The mean fluorescence spot intensity found in the single-molecule control was used as a normalization factor to convert the fluorescence intensity of *hu-yfp* and *cckA-yfp* mRNAs into the absolute copy numbers in *C. crescentus* strains, MS307 and NJH429, respectively (Fig. S5).

#### Estimated mRNA Copy Number Per Cell by smFISH

Using the copy numbers of *hu* and *cckA* mRNAs determined by smFISH, we performed a linear regression between the mRNA copy number and RPKM values of the RNA-Seq data from cells grown in M2G media^17^ (mRNA copy number = 0.0011 * RPKM R^2^=0.9952). We then used the conversion formula from the linear regression to convert the RPKM values into mRNA copy numbers across the transcriptome.

#### Estimated mRNA Copy Number Per Cell

To compare the smFISH mRNA copy numbers to independent estimates, we performed mRNA copy number estimation from RNA-seq. First, we assumed that *C. crescentus* has a similar number of mRNA per cell to *E. coli,* estimated from the literature at 1400 mRNA per cell^23^. Hence, we used this number to convert *C. crescentus* RNA-Seq relative mRNA copy number into an absolute copy number for each gene. The RPKM for *C. crescentus* RNA-Seq in log phase steady state conditions, grown in M2G media^17^ was summed across the genome, the percentage as calculated for each individual gene by dividing each gene’s RPKM by the total RPKM. Then this per gene percentage was multiplied by the total mRNA estimate for *E. coli* (1400/cell), to estimate the absolute mRNA copy number for each mRNA per cell for *Hu* and *cckA* mRNAs.

#### Individual mRNA genome-wide transcription rate

For this analysis all ORF’s half-lives were used to provide a systemic view of absolute mRNA transcription rate. Estimated mRNA copy number for each gene was divided by the decay rate to give an absolute transcription rate with in the units of molecules per cell per minute.

#### Proteomap generation

See ^32^ for explanation of how *C. crescentus* gene ontology categories were determined. For genes where a mRNA half-life had been calculated, individual gene copy number per cell was divided by the mRNA half-life in minutes to give a mRNA transcription rate in mRNA’s transcribed per minute. The categorized genes along with the mRNA transcription rate data were used to create the Proteomap^50^ for mRNA transcription rate, and mRNA half-lives were used to create the other Proteomap^50^ for mRNA half-lives.

#### 5’ Global RACE data analysis

Data was originally collected and analyzed in^28^. Since the dataset was analyzed to examine sites of 5’ PPP sites previously, we reanalyzed this dataset to search for 5’ P sites. We defined 5’ P sites as sites with < 0.594 log2 TAP ratio (Fig S3) supported with > 25 reads in both TAP+ and TAP-libraries. tRNAs which are known to be processed by RNase P to yield a 5’ P end were used as a positive control. A list of 5’ P sites fitting these criteria can be found as Table S3.

#### Sequence Logo of 5’ degradation ends

The 5’ P ends located in mRNAs from the 5’ global RACE experiment were fed into WebLogo^51^. The 5’ P site and 10nt before and after were used to generate the sequence logo.

#### RNA folding energy at 5’ P cut sites

A 50 nt sliding window was to calculate the minimum free energy by RNAfold^52^ and reported for the nucleotide centered in the middle of the window. From 100 nt upstream to 100 nt downstream of an identified 5’ P site, minimum free energy was reported as a median and graphed along with lines representing upper and lower quartiles.

#### mRNA cutting in different mRNA regions

mRNAs in this analysis were limited to those mRNA and tRNA that are monocistronic, and without multiple known isoforms (multiple TSS’s or internal TSS’s). Also, mRNAs were limited to those with known UTR’s greater than or equal to 30 nucleotides. These mRNA were divided up into different regions. With the 5’ UTR region defined as from the TSS site to –16 from CDS start. With the start region defined as –15 from CDS start to +15 from CDS start. With the coding region defined as CDS +15 from CDS start to –15 from CDS end. With the end region from –15 from CDS end to +15 from CDS end. The tRNA region was grouped together as a whole. From our 5’ end degradation detection we sorted those cuts that fell into any of the mRNA regions, or for tRNA. This gave us total number of cuts in those regions. Then the total number of cuts was divided by the length of all of those regions across our selected mRNA to give our cut site per nucleotide probability, the error bars show the calculated Wald confidence interval. Finally, the total number of cuts from the 5’ end degradation detection for each strand was randomly assigned a position in the genome, on their respective strands, using the Random Python library function “randint()”. Then the cuts that fell in each mRNA region across the genome was summed. The red line represents the total cuts in each region if cuts happened randomly across the genome as calculated in this fashion.

#### Translation efficiency vs. half-life

mRNAs in this analysis were limited to those simple mRNAs that are monocistronic, and without multiple known isoforms (multiple TSS’s or internal TSS’s) yielding 366 mRNAs. Median translation efficiency was calculated according to ^17^ mRNA were then separated into eight bins based on their translation efficiency and then the median translation efficiency of that bin was graphed vs median half-life of mRNA in that bin. Lower and upper error bars represent the 25% and 75% quartile for mRNA half-life, respectively. Center weighted ribosomal footprints Ribosomal profiling data ^32^ was used to determine the ribosomal occupancy across the genome, for footprints that had greater than 1 read per nucleotide, and the resulting footprint was center weighted. The region around the 5’ end degradation site (100 nucleotides upstream and downstream) in Fig 6C was graphed versus the ribosomal occupancy with the average set to 1 and represented by the red line. The metagene plot below shows a heat map of ribosomal occupancy (the darker red the higher ribosomal occupancy). The center represents the 5’ end degradation site, and shows where there is low ribosomal occupancy, middle ribosomal occupancy and stalled ribosomes at the degradation site.

#### Cut site probability per nucleotide

Cut site analysis was performed on highly expressed mRNAs that had greater than 1 ribosome footprint per nucleotide on average. From that set of mRNAs, the average ribosomal occupancy was calculated. Each nt along the CDS were then put into one of three categories; those nucleotides with five times or less ribosomal occupancy than average, those nucleotides with five times or more ribosomal occupancy than average, and those nucleotides with ribosomal occupancy in between. The total number of cuts in each of those ribosome occupancy categories was summed and divided by the total number of nucleotides in that category to get cuts per nucleotide probability in each category.

#### Rif-Seq with Retapamulin or Chloramphenicol pretreatment

Cell Harvesting – 25 mL of *C. crescentus* cells were grown overnight to mid-log phase in M2G media. Cells were grown in an incubation shaker, in a 250 mL conical flask at 250 rpm. Mid-log phase was confirmed as an OD_600_ reading between 0.3 to 0.5. 2 mL of Qiagen RNAprotect bacteria reagent was brought to room temperature in 15 mL falcon tubes. At time = –2 minutes (two minutes before the addition of rifampicin at time = 0 minutes) the 25 mL were pretreated with either retapamulin or chloramphenicol to a final concentration of 100 µg/mL, or retapamulin to a final concentration of 12.5 µg /mL. After this pre-incubation of retapamulin or chloramphenicol, the cells were harvested for rif-seq as described in the rif-seq section. RNA-seq libraries were deposited into NCBI GEO with accession number GSE228591. mRNA half-lives of simple mRNAs after the retapamulin and chloramphenicol treatments are listed in Table S4.

#### Codon Adaptation Index calculation

CAI was calculated across all CDSs in the *C. crescentus* genome using the CAI calculator^53^). The Codon Usage table for *C. crescentus* was obtained from Artemis gene browser.^54^. Considering the regular EYFP gene’s nucleotide sequence as the template, the codons of the sequence were randomly altered to generate sequences with different CAI values (Table S5).

#### Design of Synthetic YFP and mCherry expression plasmids

Synthetic YFP and mCherry variants with different CAI were synthesized from twist biosciences. These reporter gene fragments contained an EcoRI cut site at 5’ end and a NheI cut site at 3’ end. The pBXYFP-2 plasmid containing the kanamycin resistant gene and the reporter gene fragments were subjected to restriction digestion with EcoRI and Nhe1 and ligated together in a way that the reporter gene is under the control of an inducible xylose promoter. Inverse PCR followed by ligation was used to generate the variants with CCNA_03380 or CCNA_03024 TIR regions as described in Ghosh *et al.* 2022^16^. The list of the nucleotide sequences of reporter genes is available in the Table S5. Then the ligation mixture was transformed into E. coli cells and plated on the LB/Kanamycin plates. The resulted colonies were inoculated in 5 mL of liquid LB/Kanamycin and incubated overnight at 37 ℃ at 200 rpm. After 12 –16 hours the cultures were miniprepped using a Thermo Fisher GeneJET Plasmid Miniprep Kit. The DNA samples were sent to Genewiz for plasmid sequencing to verify the correct insert DNA sequences, using the DNA primer pXYL forward (cccacatgttagcgctaccaagtgc) and the TIR using the DNA primer UTR forward (cgtgaggccgaggatttcgcg). After the sequences were verified, the plasmids were transformed into C. crescentus NA1000 cells as described in Ghosh et al. 2022.

#### Synthetic YFP and mCherry fluorescence measurement

Fluorescence protein measurements were carried out for each of the variants using a fluorescence plate reader (Microplate Reader, Fluorescence – Gemini XPS02226). First the C. crescentus cells harboring reporter plasmids were grown overnight in liquid M2G/kanamycin medium (5 mg/mL). The next day, cells in the log phase were serially diluted with 5 mL of fresh liquid M2G/kanamycin (5 mg/mL) to have an OD_600_ of 0.05 to 0.1. The cells in the log phase were then induced with xylose (0.2 %) and the cells were grown for 6 hours at 28°C at 200 rpm. After 6 hours 150 µL of the cell culture were pipetted out to each well (three technical replicates per biological sample) of Corning 96-Well flat-bottom black microplate and fluorescence protein measurements were taken at 25℃. For YFP, exposure and emission wavelengths were 485 nm and 538 nm respectively. For mCherry, exposure and emission wavelengths were 550 nm and 620 nm respectively. The OD_600_ was measured using the same fluorescence plate reader at the time of fluorescence measurements. A *C. crescentus* strain which does not contain any fluorescence proteins (JS178) was used as the negative control. The mean fluorescence value for each variant was divided by the absorbance of the variant at 600 nm to normalize the fluorescence data. Fluorescence reading of the negative control strain was subtracted from the normalized fluorescence values of each variant to remove any background fluorescence.

#### Synthetic YFP and mCherry mRNA half-life measurement using qRT-PCR

mRNA half-life measurements were carried out for each of the variants using qRT-PCR. First the C. crescentus cells harboring reporter plasmids were grown overnight in liquid M2G/kanamycin medium (5 mg/mL). The next day, cells in the log phase were serially diluted with 25 mL of fresh liquid M2G/kanamycin (5 mg/mL) to have an OD_600_ of 0.05 to 0.1. The cells in the mid log phase (OD_600_ 0.3) were then induced with xylose (0.2 %) and the cells were grown for 2 hours at 28°C at 200 rpm. Meanwhile, 2 mL of Qiagen RNAprotect bacteria reagent is brought to room temperature in 15 mL falcon tubes. After 2 hours 1 mL of the culture was pipetted out to the 2 mL of RNAprotect and vortexed for 5 seconds to stabilize mRNAs. Then, 200 ug/mL of Rifampicin dissolved in DMSO is added to the 25 mL culture while the culture is shaking, and 1 mL of the culture was pipetted out to the 2 mL of RNAprotect and vortexed for 5 seconds at 1,3 and 9 minutes after the addition of rifampicin. After the last time point was taken, the cells were incubated at room temperature for at least 5 minutes in the RNAprotect. The cells were then spun down at 18407 x g (Eppendrof 5420, Rotor FA-24×2) for 10 minutes. The cell pellet was resuspended in 1 mL of prewarmed trizol (65°C) by gently pipetting up and down and incubated for 10 minutes at 65°C. 200 µL of chloroform was added to the mixture, gently mixed by inverting the tube few times and incubated for 5 minutes at room temperature. The sample was spun down at 18407 x g (Eppendrof 5420, Rotor FA-24×2) for 10 minutes and the supernatant was removed to a siliconized tube with 500 µl of chloroform. Subsequently, the mixture underwent vortexing and was spun for ten minutes at 18407 x g (Eppendrof 5420, Rotor FA-24×2). The supernatant was then pipetted out and combined with 700 µL of ice-cold isopropanol, 1/10 of volume of the mixture of the 3M sodium acetate and 2 µL of glyco-blue. This mixture was then incubated at –80 °C overnight. The mixture was then spun down for 60 minutes at 18878 x g (accuSpin Micro 21R Microcentrifuge, 24 x 1.5/2.0 mL Rotor) rpm at 4 °C and washed with 80 % ice-cold ethanol followed by resuspending the pelled in 30-50 µL of nuclease free distilled water. The concentration of RNA was measured using a NanoDrop 2000C from Thermo Scientific. Primers for qRT-PCR were designed using the IDT primer quest tool. Each 20 µL of qRT-PCR reaction contained Luna Universal One-Step Reaction Mix (2X), Luna WarmStart® RT Enzyme Mix (20X) Forward primer (10 µM), Reverse primer (10 µM), 200 ng of extracted RNA and nuclease free water. The reactions were quickly spun down and qRT-PCR was carried out using Applied Biosystems QuantStudio 3 Real-Time PCR System. In addition, the same qRT-PCR experiment was done using 5S forward and reverse primers to be used for the normalization of mRNA at each time point. The amount of RNA was calculated from the Ct using a standard curve. The natural log of the percentage of RNA still present in every sample was divided by the natural log of RNA at time zero. Then the slope of the linear curve fit was calculated and it was translated into mRNA half –lives by dividing –ln(2) by the slope. Primer sequences for qRT-PCR experiment are available in the Table S5.

#### Polysome profiling

##### Harvesting the cells

*C. crescentus* cells harboring reporter plasmids (CAI 0.934 and CAI 0.371 with the CCNA_03024 TIR) were grown up to mid-log phase (OD_600_ 0.3) in 200 mL of liquid M2G/kanamycin medium (5 mg/mL). Cells were then induced with xylose (0.2 %) and grown for two hours at 28°C at 200 rpm. After two hours cells were treated with chloramphenicol (final concentration of 100 μg/mL) and incubated in 28°C shaker for 2 minutes at 250 rpm. Meanwhile four 50 mL falcon tubes were filled with 15 mL ice cubes of M2G minimal media with 100 μg/mL of Chloramphenicol and the tubes were kept in ice. After 2 minutes, the cell cultures were immediately poured into 50 mL falcon tubes with the ice cubes, capped the tubes and immediately centrifuged in a prechilled centrifuge (4°C) at 22786 x g (Sorvall X1R Pro Centrifuge, F15-8 x 50cy Fixed-Angle Rotor)for 2 minutes. Then the supernatant in each tube was discarded by decanting and the cell pellets were resuspended in 20 mL of prechilled Resuspension Buffer (20 mM Tris–HCl pH 8.0, 10 mM MgCl2, 100 mM NH4Cl, 1 mM chloramphenicol). Cells were centrifuged again at 22786 x g (F15-8 x 50cy Fixed-Angle Rotor) for 1.5 minutes to pellet cells in the prechilled (rotor). Then supernatant was discarded by decanting and the cell pellet was resuspended in 600 μL of Lysis Buffer (20 mM Tris pH 8.0, 10 mM MgCl2, 100 mM NH4Cl, 5 mM CaCl2, 0.4% Triton X-100, 0.1% NP-40, 1 mM chloramphenicol, 100 U/mL RNase-free DNase I (Roche). Meanwhile, a few holes were poked in a 50mL conical lid using a hot needle and dipped in liquid nitrogen and the cell suspension in lysis buffer was slowly dripped into that 50 mL falcon tube filled with liquid nitrogen. Then the falcon tubes with the cell pellets were stored in a − 80°C freezer until proceeded with the lysis of the cells.

#### Cell lysis

Cell lysis was performed in the Mixer Mill MM 400 (Roche). First the grinding jar (25 mL) and grinding ball (25 mm) were chilled in liquid nitrogen for few minutes. Then, the jar was opened, and the frozen cell pellets were added into the chilled jar with grinding ball and the lid was closed. Then the cell pellets were milled 5 times at 15 Hz for 3 minutes. The sealed jar was chilled in liquid nitrogen between each run. Next the grinding jar was opened, and each half of the jar was kept upright in warm 30°C water for 2 minutes until the pulverized cells were thawed. Then the lysate was pipetted out into prechilled 1.5 mL microcentrifuge tube and the tubes with the Incubate lysate in ice for 20 minutes. Then the lysates were pelleted at 20,000 x g (accuSpin Micro 21R Microcentrifuge, 24 x 1.5/2.0 mL Rotor) for 10 minutes in a 4°C microfuge to remove insoluble debris and the supernatant was carefully transferred to a pre-chilled 1.5 mL microcentrifuge tube. Then 1 µL of clarified lysate was mixed with 99 µL of 10 mM Tris 7.0 and A260 of the dilution was measured by Nanodrop blanked with lysis buffer and the concentration of clarified lysate was calculated (1 A260 unit equals 40 µg/mL). Afterwards the lysates were stored in –80°C freezer until proceeded with polysome isolation using the sucrose gradient fractionation.

#### Polysome isolation by sucrose gradient fractionation

The clarified cell lysates were thawed on ice for about an hour. The SW41 rotor and buckets were chilled to 4°C. Then the open top polyclear tubes were filled with 6 mL of 55% sucrose solution and 10% sucrose solution was carefully layered on top. Then the tubes were capped and established the gradient using BIO-COMP Gradient Master. Then the caps were opened, and the cell lysates were carefully layered on the top of the gradient. The tubes were placed in buckets and the buckets were balanced. Next the buckets were spun at 35,000 rpm (209874 x g) for 2.5 hours at 4°C (Thermo Sorvall WX Ultra 80 ultra centrifuge – SW41 rotor). Then the sucrose gradients were fractionated using BIO-COMP Fractionator keeping the piston speed at 0.22 mm/s. The polysome fractions were collected to 2 mL microcentrifuge tubes and the fractions were flash frozen in liquid nitrogen and stored at –80 °C until the RNA extraction was carried out.

#### RNA extraction

To 0.7 mL of each polysome fraction, 40 µL of 20% SDS and 0.7 mL of acid phenol pre-warmed to 65°C were added and mixed by vortexing. The mixtures were Incubate at 65°C for 5 minutes and chilled on ice for 5 minutes. Tubes were spun at 20,000 x g (accuSpin Micro 21R Microcentrifuge, 24 x 1.5/2.0 mL Rotor) for 2 minutes and the aqueous phase of each fraction was transferred to fresh tubes separately. Then 0.7 mL of RT acid phenol was added, mixed by vortexing, incubated at RT for 5 minutes, spun at 20,000 x g for 2 minutes and the aqueous phase of each fraction was transferred to fresh tubes separately. Then to each tube 0.6 mL of chloroform was added, mixed by vortexing, immediately spun at 20,000 x g for 1 minute and transfer aqueous phase to fresh tubes separately. Then, 75 µL of 3 M NaOAc pH 5.5 and 0.8 mL of 100% isopropanol were added to each tube, mixed by vortexing and chill at –80°C overnight. Next day, polysomal RNA were pelleted at 20,000 x g for 1 hour in a 4°C microfuge, pellets were washed with 0.8 mL of ice cold 80% ethanol and air dried the pellets for 5 minutes and resuspended the pellets in RNase Free MQ water. Finally, qRT-PCR using Applied Biosystems QuantStudio 3 Real-Time PCR System was carried out to measure the RNA concentration in each fraction. For each qRT-PCR reaction 50 ng of RNA was used.

## Figure legends

**Figure S1: mRNA copy number comparison**.

Comparison of the mRNA copy number by mRNA FISH compared to those calculated by RNA-seq data using an assumption of 1400 mRNA/cell.

**Figure S2: Preferential cut sites at 5’ end of mRNAs**.

Metagene plot of the 5’ P site cut density across all mRNAs relative to the 5’ TSS site.

**Figure S3: 5’P Cut site identification**.

5’ end sequencing data from Zhou et al. PLOS Genetics 2015^28^ were reanalyzed to examine the site of 5’ P sites. A) Comparison of TAP treated and untreated 5’ end sequencing libraries for the tRNA^Pro^ gene. TAP ratio is displayed below, and the distribution of TAP ratios for 5’P ends of tRNAs and validated TSSs are displayed on the right. B) WebLogo^51^ of the mRNA 5’P cleavage sites. mRNA cleavage sites including 10nt before, and 10nt after the identified cleavage site were used to generate the WebLogo. C) 5’P nucleotide composition across all mRNA cut sites identified.

**Figure S4: TIR and CAI variants of mCherry show similar patterns as YFP**.

Strong (CCNA_3024) and weak (CCNA_03380) TIR regions were combined with various CAI variants of mCherry, inserted into the same pBX plasmid in NA1000 cells, and assayed for (A) mCherry fluorescence/OD600 of cells in a plate reader or (B) mRNA half-life measurement by qRT-PCR. Experiments were performed similarly to Fig 7B,C.

**Figure S5: smFISH controls**.

smFISH imaging data and quantitation for calculating single-molecule intensity. A) Measurement of FISH probe intensity for single-mRNA molecules. B) Absolute mRNA quantitation by single-molecule FISH for chromosomal HU-YFP and cckA-YFP gene fusions.

**Figure S6: smFISH probes**.

Sequences of smFISH oligonucleotide probes.

## Acknowledgements

NIGMS GM124733 to JMS. WSU Rumble fellowship to JRA. Indiana University startup funds to JMS. NSF Center for Physics of Living Cells (1430124), NSF Science and Technology Center for Quantitative Cell Biology (2243257), NIH (R35GM143203), and Searle Scholars Program to SK.

## Author Contributions

Schrader, J.M., Kim, S. Designed study, wrote paper, secured funding.

Kulathungage, H. Performed data analysis, generated all reporter constructs, performed all reporter measurements, and wrote paper.

Aretakis, J.R., Al-Husini, N., Muthunayake, N.S. Performed global Rif-seq experiments.

Aretakis, J.R. Performed data analysis and bioinformatic analysis.

Vaidya, K., Kim, S. Performed smFISH and analysis.

## Notes

### Competing Interest Statement

The authors have declared no competing interest.

