## Supplemental figures for "Rif-seq reveals *Caulobacter crescentus* mRNA decay is globally coordinated with transcription and translation"

Fig S1

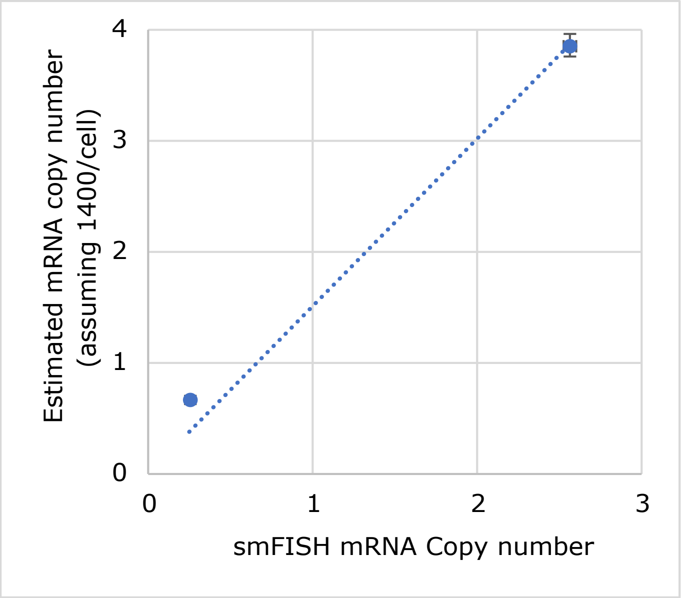
Estimated mRNA copy number
(assuming 1400/cell)
4
3
2
1
0
0
1
2
3
smFISH mRNA Copy number

Fig S2

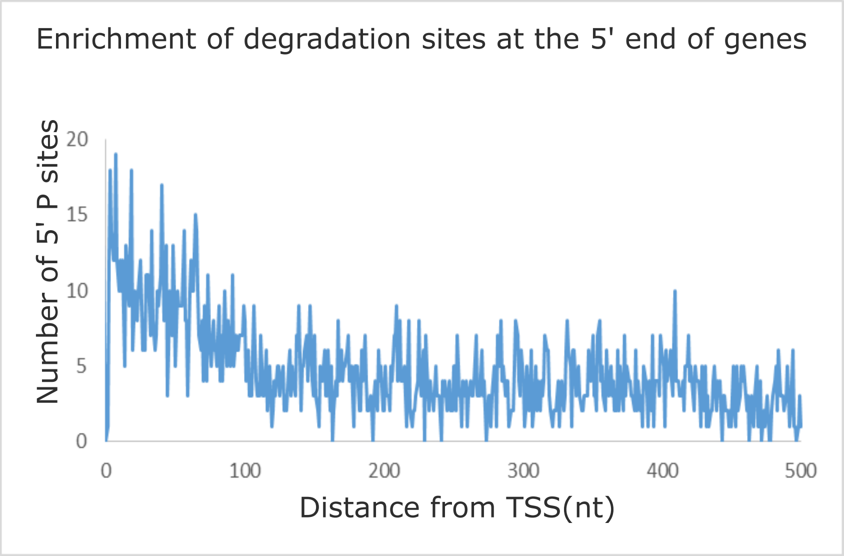
Enrichment of degradation sites at the 5' end of genes
Number of 5' P sites
20
15
10
5
0
0
100
200
300
400
500
Distance from TSS(nt)

### Fig S3

A.

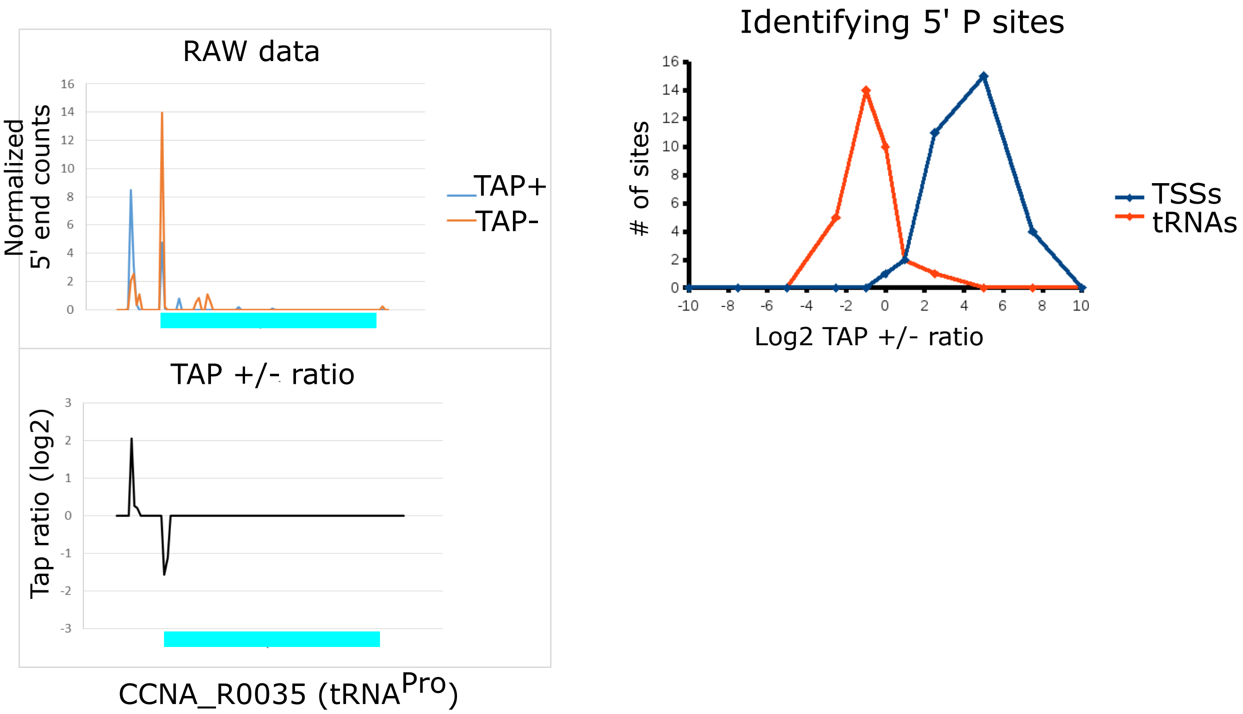
RAW data
Normalized 5' end counts
TAP+
TAP-
TAP +/- ratio
Tap ratio (log2)
CCNA_R0035 (tRNA^Pro)
Identifying 5' P sites
### of sites
Log2 TAP +/- ratio
TSSs
tRNAs

B.

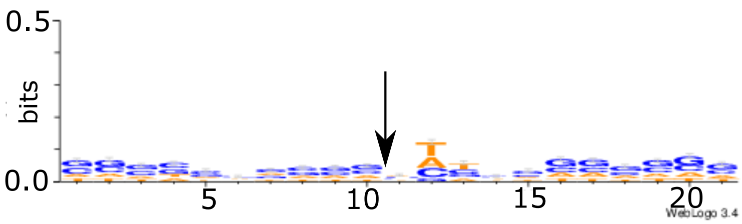
bits
0.5
0.0
5
10
15
20
WebLogo 3.4

C.

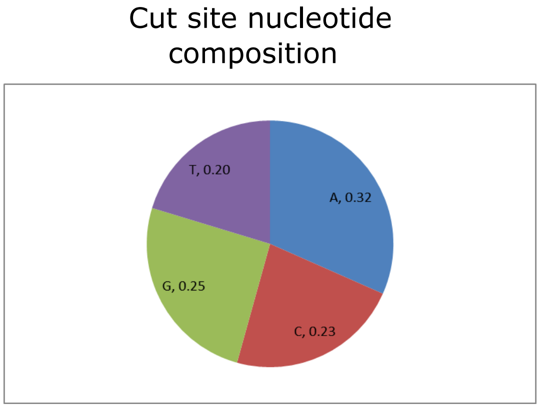
Cut site nucleotide composition
A, 0.32
C, 0.23
G, 0.25
T, 0.20

Fig S4

A.

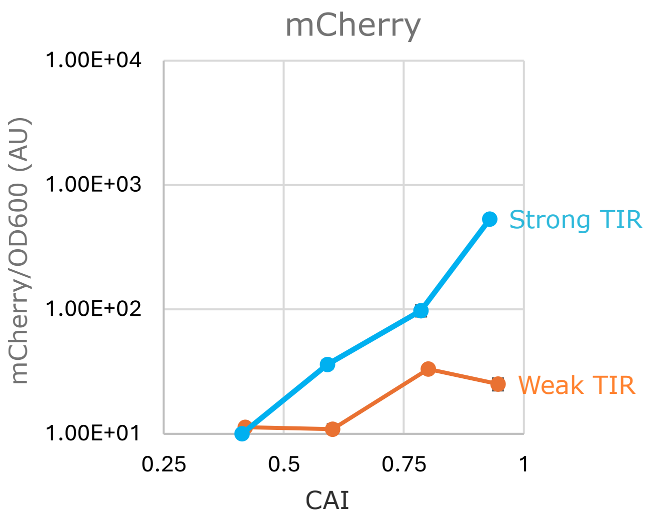
mCherry
1.00E+04
1.00E+03
1.00E+02
1.00E+01
mCherry/OD600 (AU)
Strong TIR
Weak TIR
0.25
0.5
0.75
1
CAI

B.

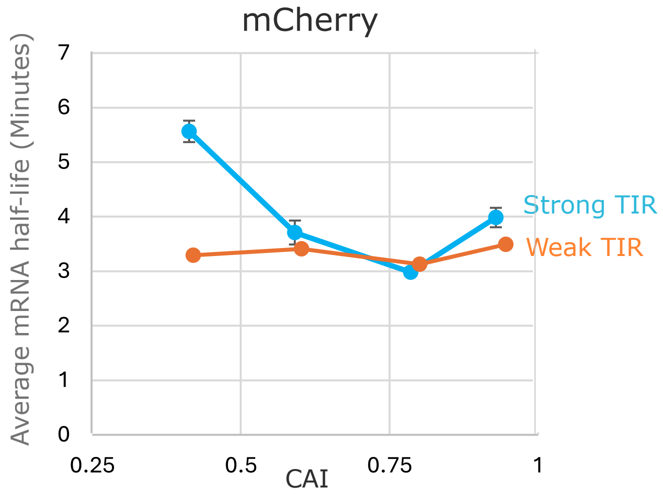
mCherry
7
6
5
4
3
2
1
0
Average mRNA half-life (Minutes)
Strong TIR
Weak TIR
0.25
0.5
0.75
1
CAI

Fig S5

A. Determining YFP single-mRNA intensity

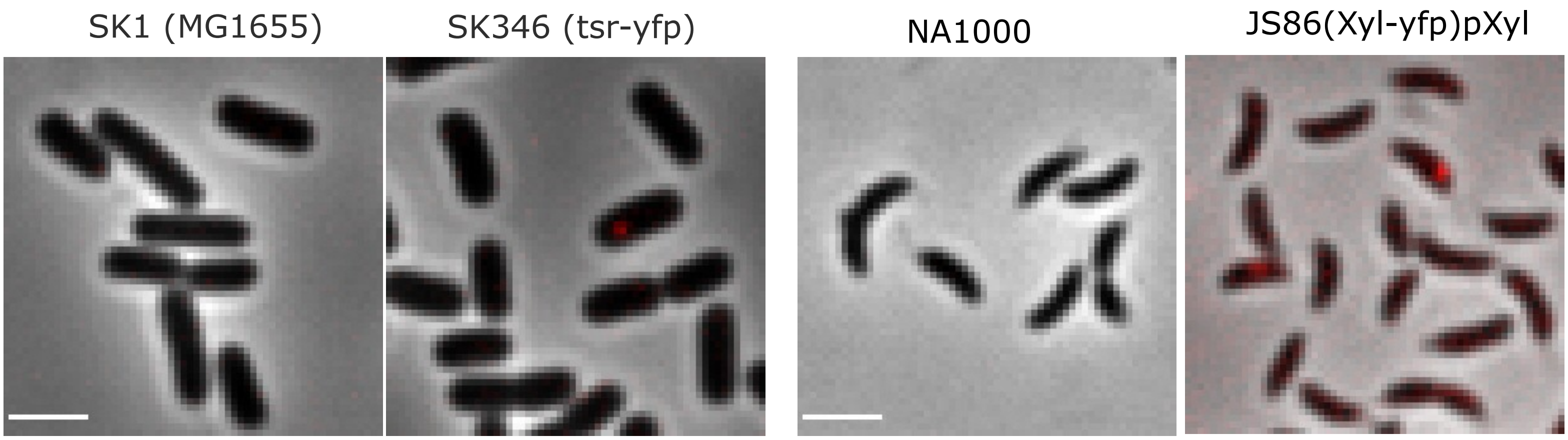


Same scale: 4300 - 10000 LUT. Scale bar = 2 micron

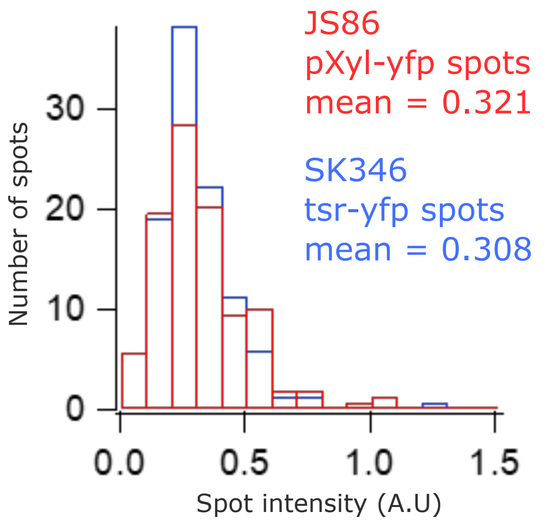


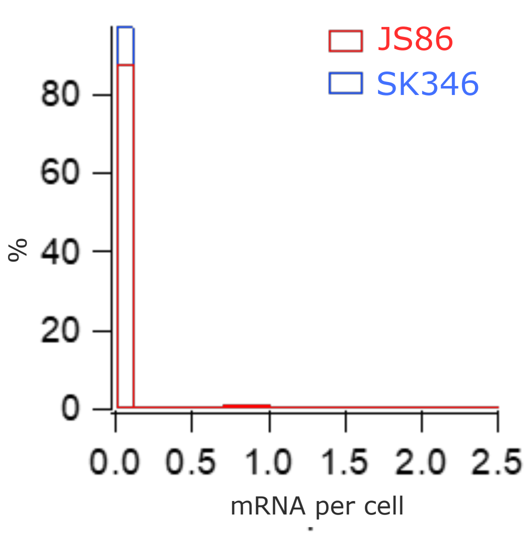


B. Absolute mRNA copy number measurement for HU-YFP and cckA-YFP mRNAs

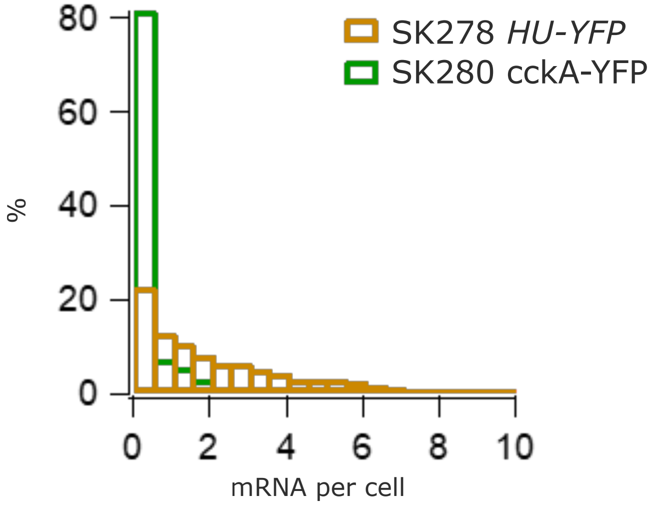

Fig S6

YFP FISH Probes

| |
|---|
| tgaacttgtggccgtttacg |
| tggtgcagatgaacttcagg |
| tagccgaaggtggtcacgag |
| tggcggacttgaagaagtcg |
| cttgaagaagatggtgcgct |
| ttgaagtcgatgcccttcag |
| ctgttgtagttgtactccag |
| ttgtcggccatgatatagac |
| cttgaagttcaccttgatgc |
| tagctcaggtagtggttgtc |
| gtcacgaactccagcaggac |
| tacttgtacagctcgtccat |
